# Circadian Dysregulation in Aging Alters Senescence and Inflammatory Pathways in a Sex- and Time-of-Day–Dependent Manner

**DOI:** 10.64898/2026.03.05.709919

**Authors:** Gretchen T. Clark, Yue Zhao, Robyn E. Reeve, Colleen M. Farley, Courtney Willey, Susan Sheehan, Samantha Spellacy, Alicia Warren, Abigail Brackett, Nadia A. Rosenthal, Ron Korstanje

## Abstract

The circadian rhythm orchestrates gene expression and critical physiological processes but becomes disrupted with aging, contributing to disease. How this disruption interacts with cellular senescence—a key driver of aging pathology—remains poorly defined. We studied renal gene expression at four timepoints over 24hrs in 6- and 24-month-old genetically diverse UM-HET3 mice of both sexes and performed complementary analyses in synchronized fibroblasts sampled at seven timepoints. Aging dysregulated core clock relationships, including loss of the canonical anti-phase expression between *Bmal1* and *Per2*. Senescence-associated genes were not static but exhibited pronounced oscillations, with senescence phenotypes varying by sex and time of day. Differential expression analysis revealed immune activation, metabolic rewiring, and epigenetic changes that were sex- and time-dependent. Variance analysis uncovered increased transcriptional noise in aging, particularly in circadian-regulated pathways such as RNA splicing, ribosome biogenesis, and TOR signaling. Single-nucleus RNA-Seq identified two cell populations lacking the normal *Bmal1*–*Cdkn1a* expression relationship: one senescent-like and another profibrotic, revealing distinct cell states linked to circadian dysregulation. Fibroblasts recapitulated key age-related circadian changes seen in the kidneys, including phase shifts in mTOR and oxidative phosphorylation. Together, this work demonstrates that senescence phenotypes are dynamic, sex-specific, and time-of-day dependent, and introduces a new framework for detecting senescent cells based on circadian gene relationships. These findings underscore the need to integrate temporal context into aging research and therapeutic strategies.

**Graphical Abstract:** 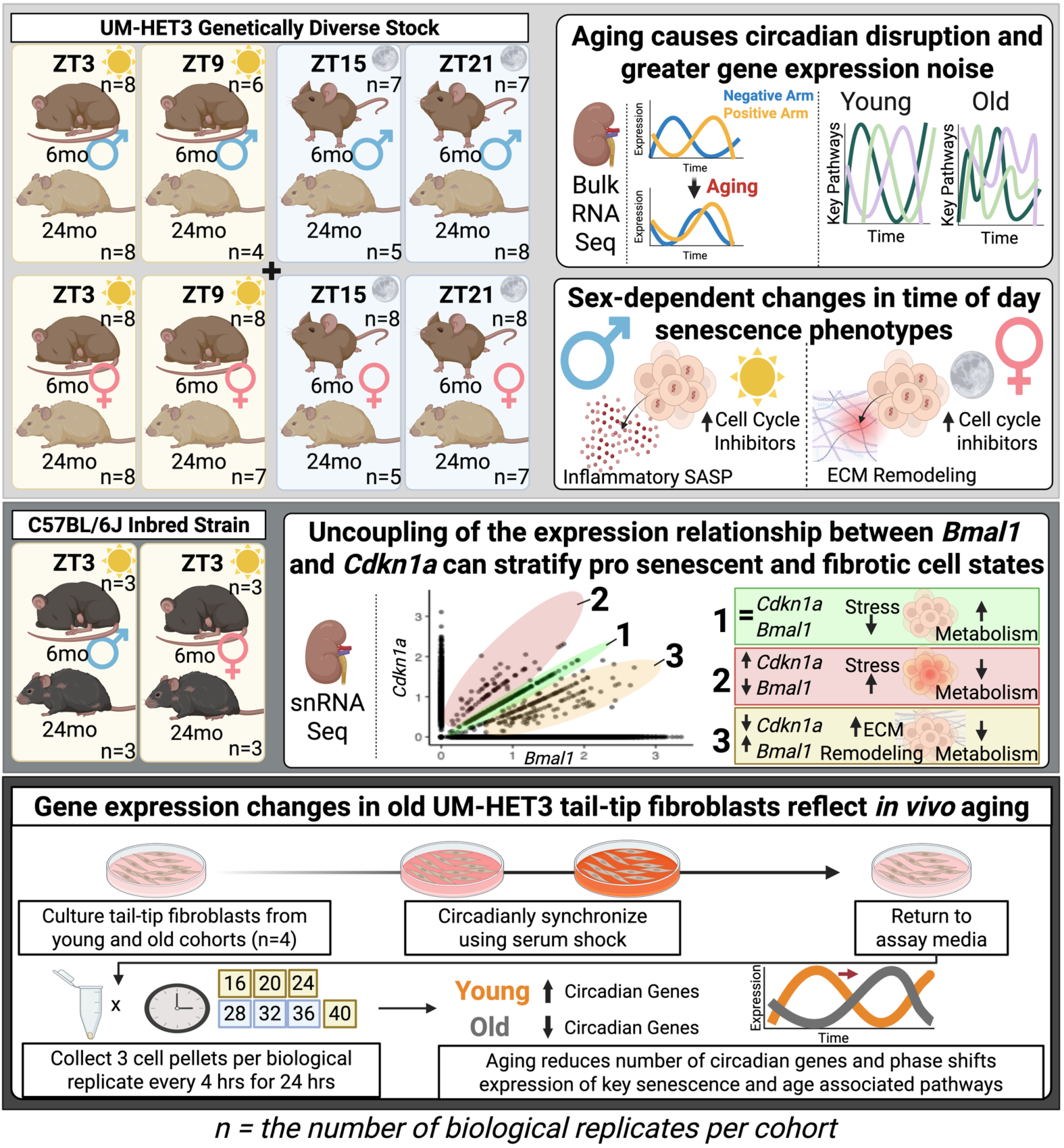

## 1. Introduction

Circadian rhythms are intrinsic 24-hour oscillations that synchronize physiology with environmental light–dark cycles, coordinating processes such as metabolism, cell cycle progression, and stress responses (Collins et al., 2021; García Cobarro, Ignez Soares, Kutsenko, & Tomas-Loba, 2025; Matsu-Ura et al., 2016). These rhythms are driven by a transcription–translation feedback loop initiated when the positive arm transcription factors BMAL1 and CLOCK heterodimerize and bind E-box elements to activate clock-controlled genes, including their own repressors, Period (*Per1, Per2, Per3*) and Cryptochrome (*Cry1, Cry2*). PER and CRY proteins accumulate in the cytoplasm, undergo sequential phosphorylation, and form inhibitory complexes that translocate to the nucleus to suppress BMAL1/CLOCK activity. This repression persists until the complex is hyperphosphorylated, ubiquitinated, and degraded, restarting the cycle (Partch, Green, & Takahashi, 2014). Two auxiliary loops stabilize this core oscillator: the RORα/REV-ERBα loop, which regulates *Bmal1* transcription, and the NFIL3/DBP loop, which modulates expression of *Per* and other clock-controlled genes (Partch et al., 2014; Yoshitane et al., 2019). Together, these interlocking loops maintain temporal homeostasis and synchronize cellular processes with external cues.

With aging, the circadian system often becomes disrupted, manifesting as altered amplitude, phase, and robustness of rhythmic gene expression (García Cobarro et al., 2025). Such disruption has been implicated in the pathogenesis of age-related diseases, including cancer, cardiovascular disease, neurodegenerative disease, and metabolic disorders (García Cobarro et al., 2025). Mechanistically, aging disrupts pathways under circadian control—such as mTOR signaling, oxidative phosphorylation, and DNA repair—while impairing nutrient sensing and mitochondrial function (Cao, 2018; Collins et al., 2021; García Cobarro et al., 2025). These same processes are central to cellular senescence, a state of irreversible growth arrest characterized by chromatin remodeling, metabolic reprogramming, and secretion of pro-inflammatory factors collectively termed the senescence-associated secretory phenotype (SASP) (Suryadevara et al., 2024). Senescence can be triggered by DNA damage, replication stress, and mitochondrial dysfunction—all processes that intersect with circadian regulation (García Cobarro et al., 2025). Despite these connections, the temporal dynamics of senescence in aging tissues remain poorly defined, and inconsistent identification of senescence markers across studies suggests that circadian oscillations may underlie this variability (Cohn, Gasek, Kuchel, & Xu, 2023).

If senescence phenotypes oscillate throughout the day, then sampling at a single timepoint could obscure or misrepresent their molecular signatures, contributing to conflicting reports. Furthermore, sex differences in circadian regulation and aging are well documented, yet their combined impact on senescence has not been systematically explored (Paschos, Lordan, & FitzGerald, 2025). Studies in inbred mouse strains further limit generalizability, leaving open questions about how these interactions manifest in genetically diverse populations that better model human heterogeneity. Most aging studies describe circadian changes broadly as circadian disruption, meaning systemic misalignment between internal rhythms and external cues. Because the circadian system relies on precise timing between its positive and negative arms, we hypothesized that aging alters these internal relationships—a form of circadian dysregulation that affects phase coordination even when oscillations persist. We further reasoned that this internal degradation may extend downstream, weakening the timing between the clock and the pathways it regulates. We term this loss of coordinated timing circadian uncoupling. These distinctions guided our examination of how aging reshapes internal circadian timing.

To address these gaps, we characterized circadian transcriptomes in kidneys and tail-tip fibroblasts from genetically diverse UM-HET3 mice at 6 and 24 months of age. UM-HET3 mice originate from a four-way cross of C3H/HeJ, BALB/cByJ, C57BL/6J, and DBA/2J, producing broad genetic heterogeneity comparable to that in humans (Jackson, Fornés, Galecki, Miller, & Burke, 1999). Bulk RNA-seq captured genome-wide circadian oscillations and age-related expression changes in kidney tissue. Because bulk profiles obscure cell-type heterogeneity, we complemented this approach with single-nucleus RNA-seq (snRNA-seq) in C57BL/6J mice to resolve cell specific changes in clock–senescence interactions. Finally, phase-set enrichment analysis (PSEA) of tail-tip fibroblast transcriptomes revealed age-dependent phase shifts in key pathways, closely mirroring patterns observed in the kidney.

By integrating time-of-day, sex, and genetic diversity, this study provides mechanistic insight into clock–senescence interactions, explains variability in senescence marker detection, and introduces a relational framework for identifying senescent states based on circadian–senescence gene uncoupling.

## 2. Results

### 2.1 Aging causes renal circadian dysregulation in both male and female genetically diverse mice

We hypothesized that aging degrades the expression relationships between clock genes, resulting in circadian dysregulation. To assess this, we collected UM-HET3 kidneys at 6 and 24 months every six hrs over 24hrs, sampling at four Zeitgeber timepoints (ZT3, ZT9, ZT15, ZT21). We profiled gene expression by performing bulk RNA-seq on whole-kidney tissue and aligned reads using the Genome Reconstruction by RNA-seq (GBRS) pipeline (Choi et al., 2025). Gene expression was then normalized to transcripts per million (TPM) for all downstream circadian analyses. Using Generalized Linear Mixed Modeling (GLMM) Cosinor modeling with Diagnostics for Hierarchical Regression Models (DHARMa) residual checks, we identified rhythmic genes in all cohorts and confirmed that many canonical clock components—such as *Bmal1, Clock, Per*, and *Cry*—were classified as circadian in both young and old kidneys (Figure S1, Tables S1 and S2). These findings suggest that core clock components remain rhythmic with age, though interpretation is constrained by DHARMa residuals and model overfitting which prevented several genes from being called oscillatory in young samples. This may partly reflect the limited timepoint resolution, though the dataset spans 24hrs. Thus, the presence or absence of rhythmicity alone cannot fully capture circadian health in this dataset.

To probe organization of the clock, we next examined the expected anti-phase relationship between the activator-driven positive arm and the repressor-driven negative arm. Using TPM-normalized kidney RNA-seq data from all cohorts, we plotted normalized (0–1) expression values for each core clock gene during timepoints when expression exceeded the gene’s mean expression across the cycle (Figures 1A–1D). Young mice displayed clear separation between the expression of positive and negative arms with the negative arm peaking during the daytime (ZT3, ZT9) and the positive arm during the night (ZT15, ZT21). In contrast, aging altered these dynamics: in males, old animals lost the daytime peak of *Csnk1e* and showed a shortened daytime expression window for *Nr1d1* (*Reverbα*) (Figures 1A and 1B). In females, negative-arm changes were more pronounced, with loss of daytime peak expression for *Csnk1e*, *Per2*, and *Per1* (Figures 1C and 1D).

**Figure 1.**
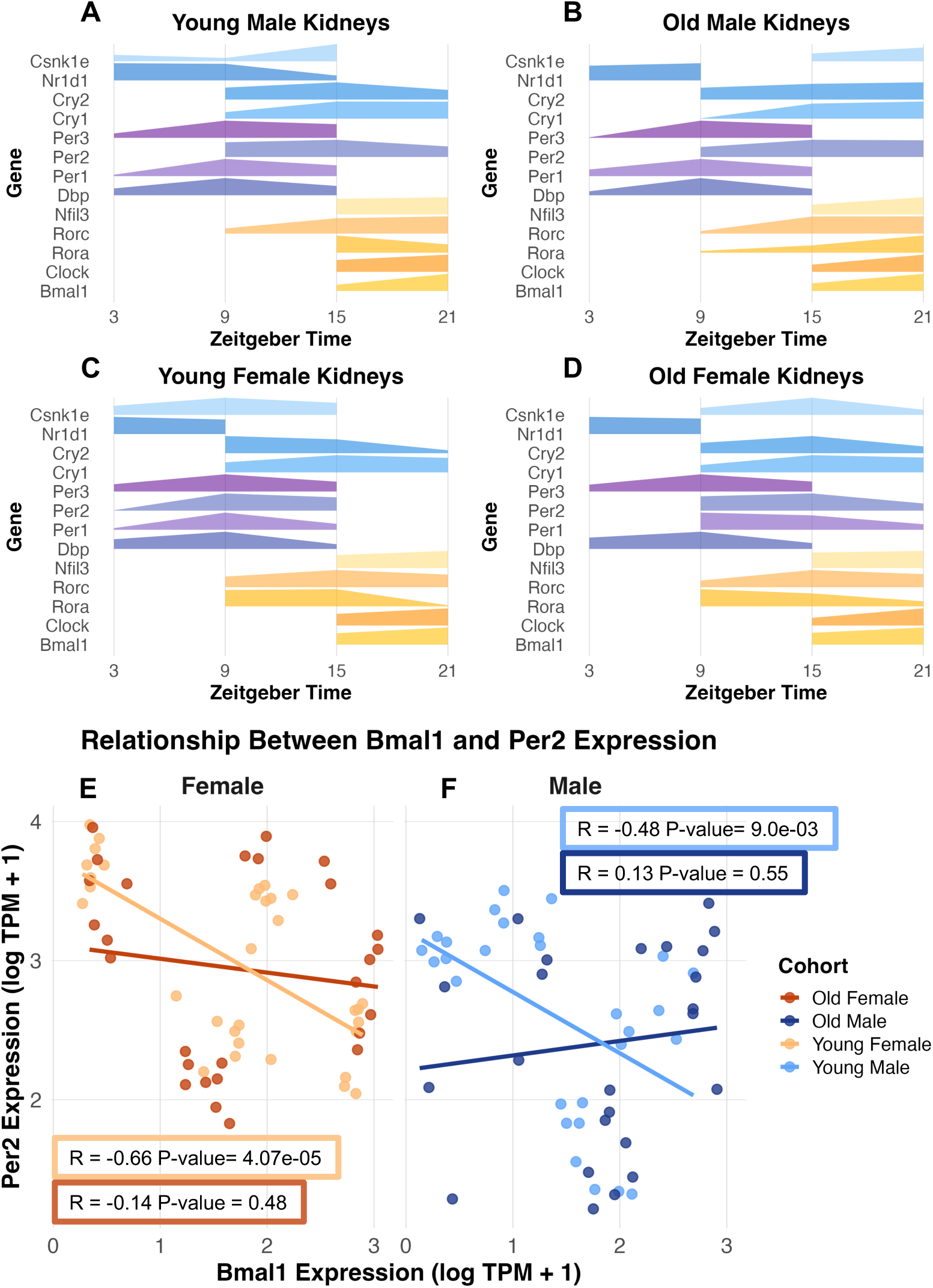
Age-related alterations in circadian gene expression in kidneys. A–D). Phase plots showing genes with expression above the mean over the circadian cycle with increased line thickness showing higher expression for young males (A), old males (B), young females (C), and old females (D). Negative-arm components are shaded in blues and purples, and positive-arm components in oranges and yellows. E). Correlation between *Bmal1* and *Per2* expression across ZT in young (light orange) and old (dark orange) female kidneys plotted as log(TPM + 1). F). Correlation between *Bmal1* and *Per2* expression across ZT in young (light blue) and old (dark blue) male kidneys plotted as log(TPM + 1).

To further quantify these changes, we looked at the correlation of expression of key clock genes, *Bmal1* and *Per2*. In young kidneys, these genes exhibited a strong negative correlation (young males, R = -0.48, p-value = 9.0e-03; young females, R = -0.66, p-value = 4.07e-05), consistent with an antiphase expression (Figures 1E and 1F). In contrast, this relationship was lost in both sexes with age (old males, R = 0.13, p-value = 0.55; old females, R = -0.14, p-value = 0.48), indicating a breakdown in temporal coordination despite residual rhythmicity (Figures 1E and 1F). This dysregulation suggests that aging does not simply dampen oscillations but alters the underlying network architecture, potentially impairing downstream processes such as metabolism, stress responses, and senescence.

Together, these results establish that aging erodes circadian integrity in genetically diverse mice. Importantly, using relational metrics—such as the anti-phase coupling of *Bmal1* and *Per2*—allowed us to detect this dysregulation even with the limited temporal resolution of four timepoints, highlighting the value of expression relationships with these limitations.

### 2.2 Time of day affects senescence phenotypes in aged kidneys

Next, we investigated if time of day affects senescence gene expression. Senescence is a hallmark of aging, yet its molecular signature has been inconsistent across studies and tissues (Cohn et al., 2023; Lopez-Otin, Blasco, Partridge, Serrano, & Kroemer, 2023). We hypothesized that this variability reflects circadian regulation of senescence-associated genes, causing time-of-day–dependent changes in their expression. To test this, we applied GLMMcosinor modeling with DHARMa residual checks to canonical senescence markers and the SenMayo gene set (Saul et al., 2022). Genes were classified as circadian if they exhibited significant sine or cosine oscillations, amplitude, phase estimates, and passed DHARMa residual checks for model fit (Tables S3 and S4).

Our analysis revealed that senescence phenotypes are not static but oscillate in expression over the day with aging amplifying these dynamics. We found that aging does not uniformly increase senescence markers. Instead, it introduces new rhythmic components while shifting the timing of existing ones, expanding circadian control over the senescence program. For example, *Cdkn1a*, *Ccl5*, *Mif*, *Il15*, *Igfbp4*, and *Serpine1* were circadian in all cohorts, while others changed status with age: *Igfbp3* and *Plaur* were rhythmic only in young kidneys, whereas *Ctnnb1*, *Cxcl10*, *Iqgap2*, *Mmp3*, and *Selplg* gained rhythmicity in old kidneys (Figure S2, Tables S3 and S4). These genes regulate signaling, immune chemotaxis, adhesion, and extracellular matrix (ECM) remodeling—processes central to SASP evolution and tissue remodeling in aging (Saul et al., 2022).

Differential expression analysis (DESeq2; adjusted p-value < 0.05) comparing young and old samples within each sex and across timepoints confirmed that SASP profiles vary by both sex and time of day (Figures 2A and 2B). Early light phase (ZT3) marked SASP initiation in both sexes, with males expressing inflammatory cytokines/chemokines (*Ccl2/5/7, Cxcl1/2/10, Csf1*) and females expressing inflammatory-ECM remodeling genes (*Mmp3, Selplg, Serpine2, Icam1*). In the late light phase (ZT9), males intensified cytokine/chemokine-driven inflammation (*Cxcl2/16*, *C3*), while females expanded pro-inflammatory protease programs (*Mmp3, Mmp9, Selplg, Serpine2, Mmp14, Icam1*). By early night (ZT15), females exhibited the most extensive SASP signature, including complement activation (*C3, Cd55↓*), growth factor modulation (*Igfbp4/5/7*), and cytokines/chemokines (*Ccl/5/8, Csf1, Mif*). Males appeared largely quiescent at this timepoint, but this likely reflects reduced statistical power due to fewer aged male replicates rather than true biological absence of SASP activity. Late dark phase (ZT21) saw females with the smallest proinflammatory (*C3, Csf2rb, Cd55↓*) and ECM remodeling signature (*Serpine1/2, Icam1*), while males shifted toward mixed inflammation (*C3, Ccl2, Il17, Csf1)* and growth factor signaling (*Egf, Hgf, Nrg1, Angpt1, Bmp6*).

**Figure 2.**
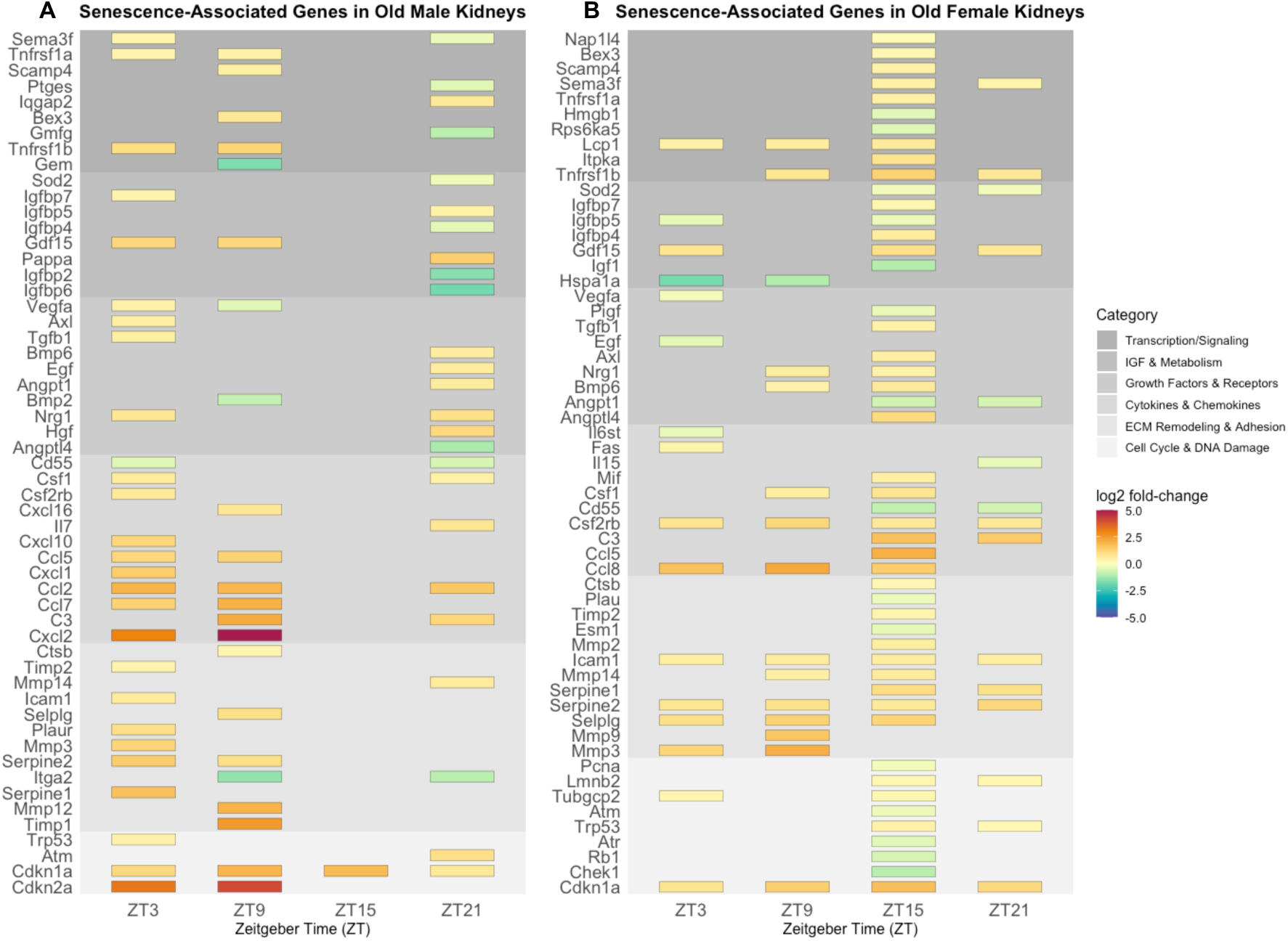
Time-of-day and sex-dependent differences in senescence-associated gene expression. A). Significantly (padj < 0.05) differentially expressed genes in old male kidneys compared to young male kidneys over Zeitgeber Time (ZT). B). Significantly (padj < 0.05) differentially expressed genes in old female kidneys compared to young female kidneys over ZT. Genes are categorized based on their known function in senescent cells. Upregulated genes are shown in red and orange and downregulated genes are shown in green and blue.

Together, these findings show that senescence signatures are dynamic, sex-specific, and strongly time-of-day dependent. Males display a daytime inflammatory senescence phenotype (ZT3, ZT9) that diminishes at night, whereas females show increasing inflammatory and ECM remodeling signatures from day to night (ZT9 to ZT15). Additionally, we saw sex difference in the differential expression of key senescence and age-related gene, *Cdkn2a* which was upregulated in old males but only during day timepoints and was not upregulated in old females at any time point (Figures 2A and 2B). In contrast, *Cdkn1a* was consistently upregulated with aging in both sexes across all timepoints (Figures 2A, 2B, S2), indicating its reliability as a marker of aging across sex and time.

### 2.3 Aging alters renal immune and metabolic pathways in a sex- and time-dependent pattern

We next examined how aging alters biological pathways across circadian timepoints, focusing on enriched gene ontologies that capture known aging signatures such as increased inflammation, impaired repair, and metabolic disruption (Lopez-Otin et al., 2023). To characterize pathway-level changes, kidneys from young and old UM-HET3 mice were compared within each sex at ZT3, ZT9, ZT15, and ZT21 using DESeq2 for differential expression analysis (adjusted p < 0.1; log₂FC ≥ 1 or ≤ −0.5). Gene Ontology enrichment was performed with clusterProfiler on up and down regulated genes separately, and the significant terms (ranked by adjusted *p*-value) were summarized by computing semantic similarity using rrvgo with a similarity threshold of 0.5 to reduce redundancy.

The early light phase (ZT3) showed immune activation in both sexes, with upregulation of mononuclear and lymphocyte proliferation, leukocyte expansion, and humoral responses (Figures 3A and 3B). This shared signature persisted at ZT9, where males continued to enrich leukocyte, lymphocyte, and myeloid proliferation pathways, and females maintained robust B-cell activation and immune-signaling programs (Figures 3A and 3B). In the dark phase (ZT15–ZT21), the sexes diverged more clearly. Males shifted toward metabolic activation, including fatty acid metabolism and chemokine signaling, whereas females sustained broad immune enrichment (Figures 3A and 3B). By the late dark phase (ZT21), males further increased lipid localization, lipid transport, stimulus-response pathways, and humoral immunity, while females remained enriched for immune ontologies such as somatic recombination, complement activation, and B-cell–mediated immunity (Figures 3A and 3B). Overall, females maintained immune activation across the circadian cycle, whereas males displayed a daytime immune peak followed by nighttime activation of lipid-related metabolic programs, which aligns with the daytime peak of SASP-associated inflammation (Figure 2A).

**Figure 3.**
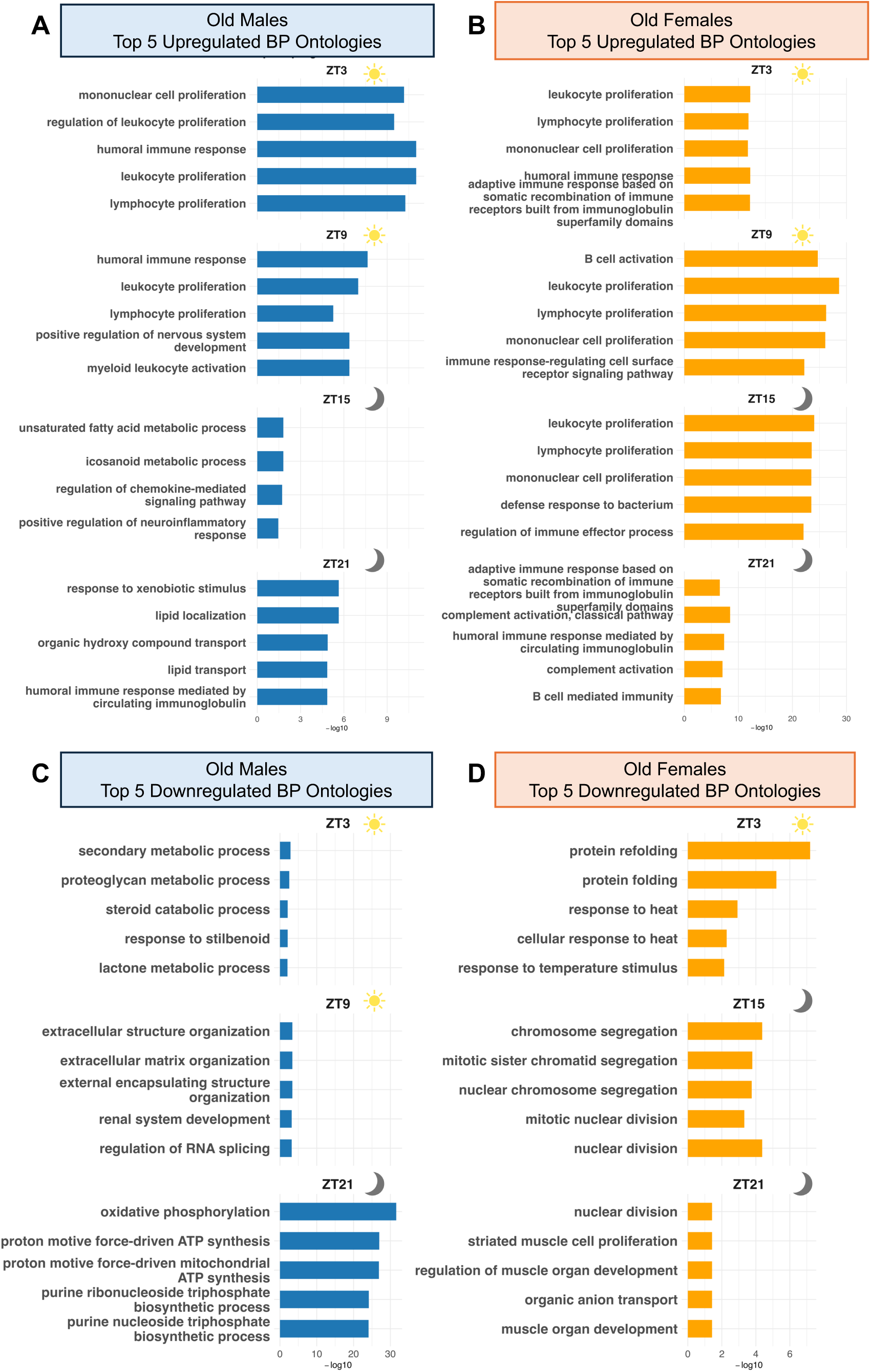
Time-of-day and sex-dependent alterations in gene ontologies in old kidneys. A–B). Bar plots displaying the top five upregulated Biological Process ontologies in old male (A) and old female (B) kidneys relative to young counterparts across four Zeitgeber Time (ZT) points. C–D). Bar plots displaying the top five downregulated Biological Process ontologies in old male (C) and old female (D) kidneys across the same ZT points. Enrichment significance is represented as –log₁₀(FDR). Sun icons denote daytime ZT intervals, while moon icons denote nighttime ZT intervals.

Downregulated pathways showed similarly distinct sex-specific timing. At ZT3, males suppressed proteoglycan- and steroid-related metabolic processes, while females downregulated protein folding and heat-response pathways (Figures 3C and 3D). By ZT9, males further downregulated ECM organization, renal development, and RNA splicing, whereas females showed minimal repression limited to lipid and carboxylic acid transport (Figures 3C and S3). Divergence increased at ZT15: males downregulated histone H3K4 trimethyltransferase (Figure S3), whereas females repressed multiple cell-cycle ontologies, coinciding with elevated SASP expression (Figures 3D and 2B). By ZT21, males exhibited downregulation of oxidative phosphorylation and purine biosynthesis, indicating potential mitochondrial and energy imbalance, while females suppressed additional cell-cycle and muscle-development pathways (Figures 3C and 3D).

These findings reveal that aging-driven immune activation and metabolic remodeling are not static but vary by sex and time of day. Notably, females display more persistent and broad immune enrichment across timepoints, whereas males show greater temporal shifts from immune to metabolic programs, underscoring distinct trajectories of aging between sexes.

### 2.4 Aging kidneys increase variability in expression of genes linked to major aging pathways

We next examined gene expression variability in aging, a key hallmark of age-related decline that reflects increasing stochastic dysregulation of pathways maintaining cellular homeostasis (Lopez-Otin et al., 2023; Wolff et al., 2023). By sampling kidneys at multiple timepoints (ZT3, ZT9, ZT15, and ZT21), we could visualize dynamic rhythmic fluctuations that single-timepoint studies often overlook. We hypothesized that circadian sampling would uncover variability in pathways under temporal control, providing a more complete picture of transcriptional instability in aging.

Variance analysis was performed using missMethyl on RNA-seq data from young and old UM-HET3 kidneys at all timepoints. Raw counts were normalized using voom transformation to model the mean–variance relationship, and duplicate correlation was applied to account for repeated circadian measurements. Differential variability (DV) was assessed using Levene’s test-based models, and genes were considered significant at an adjusted *p*-value < 0.05. Gene Ontology enrichment of DV-associated genes was performed using clusterProfiler, and redundancy among terms was reduced using rrvgo clustering with a similarity threshold of 0.9 and a dispensability cutoff < 0.1. From these clusters, the top 20 representative pathways ranked by adjusted p-value were selected for visualization and interpretation (Figures 4A and 4B).

**Figure 4.**
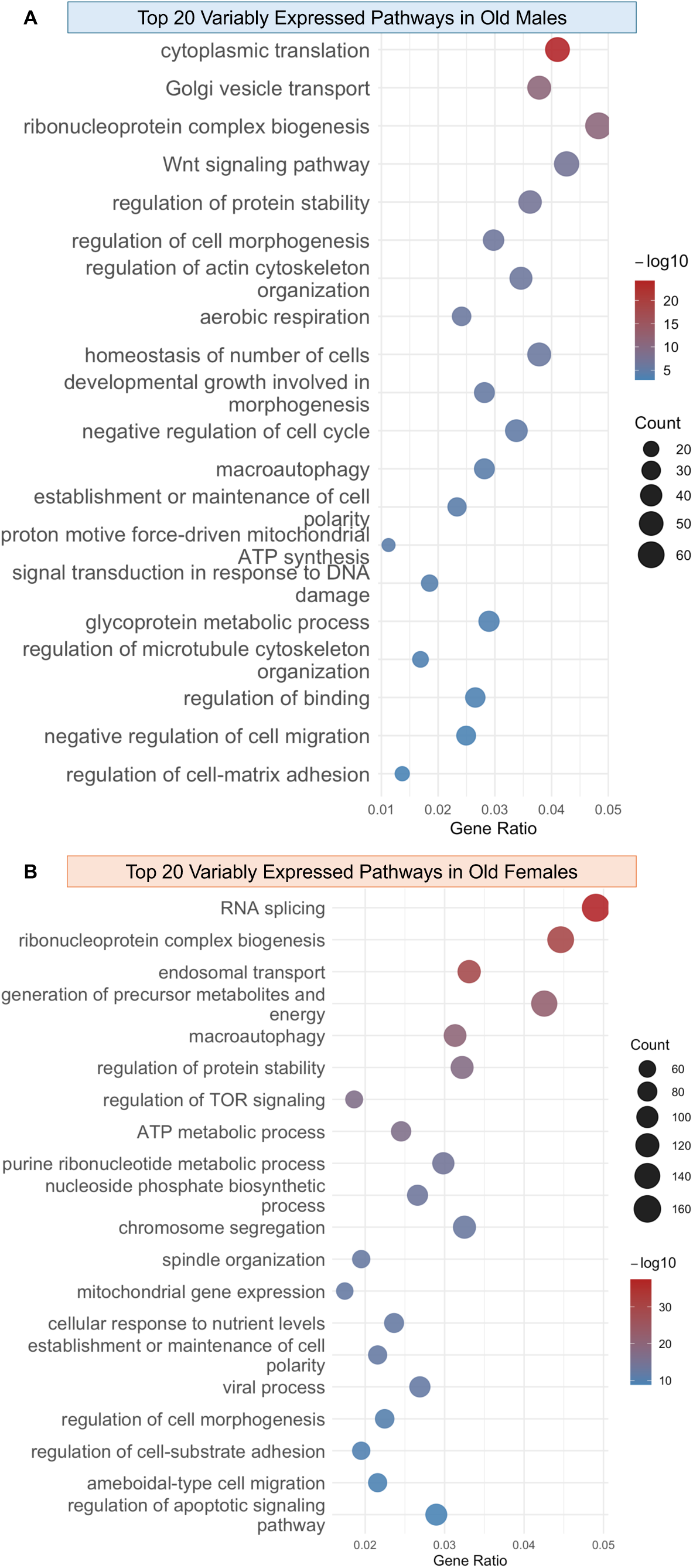
Aging kidneys have increased variability in the expression of key pathways. Bubble plots of the top 20 most variably expressed pathways ranked by false discovery rate (FDR): (A) aged males, (B) aged females. Bubble size corresponds to the number of genes in the pathway; color indicates -log of q values of the pathways, with red representing higher significance and blue representing lower significance.

Our analysis revealed a striking increase in transcriptional variability with age, particularly in pathways central to proteostasis, energy metabolism, and stress response. In aged males, highly variable pathways included cytoplasmic translation, Golgi vesicle transport, ribosome biogenesis, Wnt signaling, and negative regulation of the cell cycle, indicating instability in protein synthesis and signaling control (Figure 4A). Females exhibited variability in RNA splicing, macroautophagy, TOR signaling, ATP metabolism, and chromosome segregation, pointing to disrupted RNA processing and nutrient-sensing pathways (Figure 4B). When sexes were combined, the top enriched pathways included RNA splicing, ribosome biogenesis, macroautophagy, TOR signaling, ATP synthesis, and oxidative stress response—core processes consistently implicated in aging biology (Figure S4) (Lopez-Otin et al., 2023). Notably, variability extended beyond stress pathways to core processes such as translation initiation, organelle organization, and cell cycle regulation, suggesting widespread transcriptional instability.

Critically, many of these pathways are themselves circadianly regulated such as translation, ribosome biogenesis, Wnt signaling, cell cycle, RNA splicing, mTOR signaling, and ATP synthesis meaning that static sampling would have underestimated their variability (Cao, 2018; Collins et al., 2021; García Cobarro et al., 2025; Matsu-Ura et al., 2016; McGlincy et al., 2012; Partch et al., 2014). By capturing multiple timepoints, we revealed that aging amplifies transcriptional noise in time-sensitive processes, disrupting rhythmic control of translation, cell cycle, and nutrient sensing. This finding reframes variability as a dynamic, circadian-linked feature of aging rather than random noise, and demonstrates that without temporal resolution, key signatures of aging biology remain invisible.

### 2.5 The uncoupling between *Cdkn1a* and *Bmal1* expression can stratify senescent-like and profibrotic cell phenotypes in snRNA-seq data

Given our findings of circadian dysregulation in aging and time-of-day variability of senescence marker expression, we hypothesized that circadian uncoupling could underly senescence progression. To investigate this, we quantified the relationship between the circadian activator and known cell cycle regulator *Bmal1* and the senescence effector *Cdkn1a* in bulk kidney RNA-seq data across all four Zeitgeber timepoints (ZT3, ZT9, ZT15, ZT21) (García Cobarro et al., 2025; Suryadevara et al., 2024). In young kidneys, there was a strong linear correlation between *Bmal1* and *Cdkn1a* expression (young males, R = 0.914, p-value = 1.07e-11; young females R = 0.729, p-value = 2.25e-06) (Figures 5A, light orange and 5B, light blue). With age, this linear relationship weakened in both sexes: old males retained a correlation of 0.804 (p-value = 1.29e-06) while old females dropped to 0.680 (p-value 9.51e-05) (Figures 5A, dark orange and 5B, dark blue). Interestingly, the y-intercept of the correlation increased with age (males: 0.924→2.064; females: 1.30→2.012), indicating an elevated baseline of *Cdkn1a* expression. Together, these data show that aging does not abolish expression of either gene but erodes the strength of the *Bmal1–Cdkn1a* relationship and lowers their correlation, with a marked increase in baseline *Cdkn1a* expression. This phase-coupling metric between *Bmal1* and *Cdkn1a* may therefore provide a more sensitive indicator of cellular stress or emerging senescence than either marker alone.

**Figure 5.**
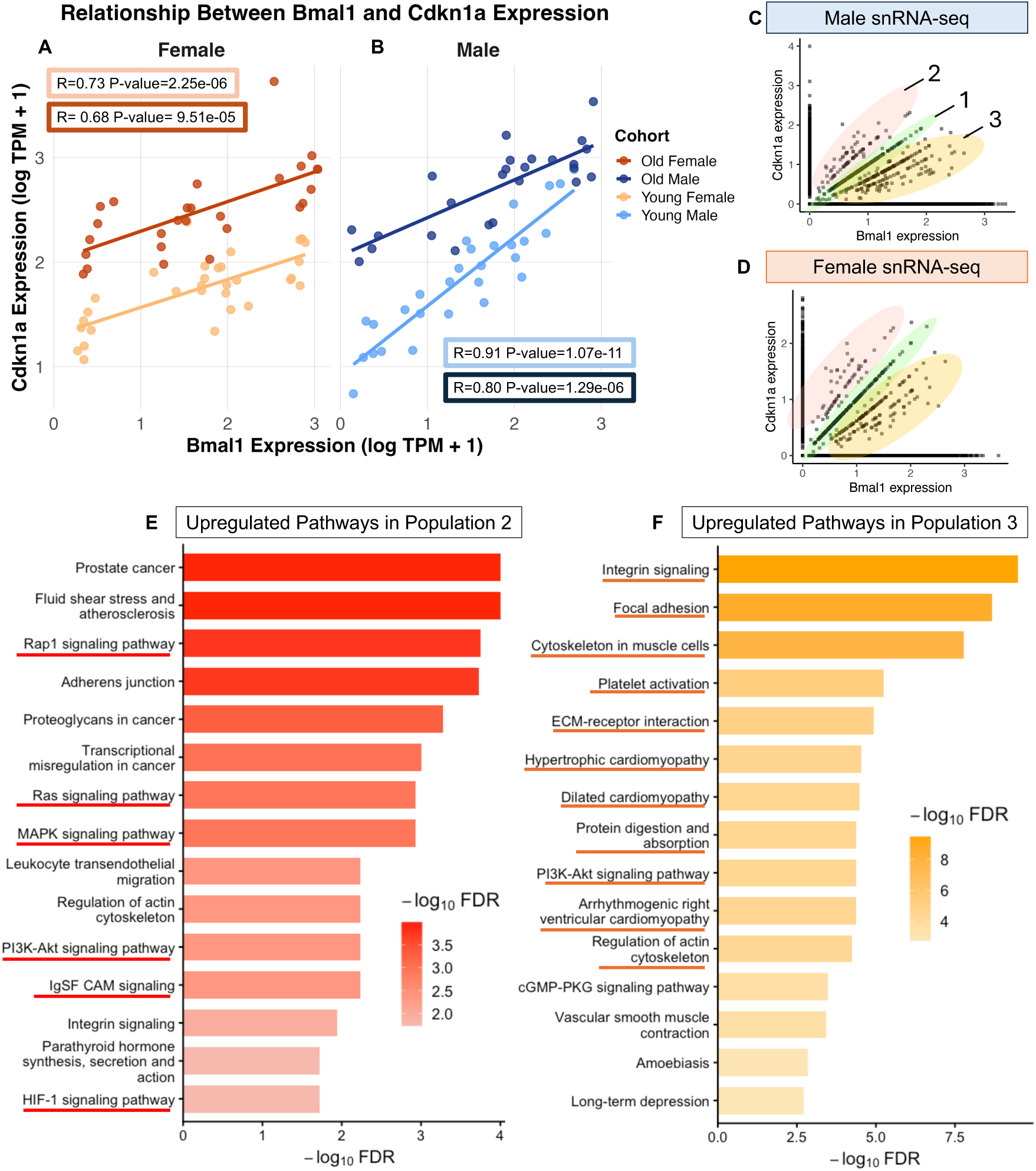
*Bmal1–Cdkn1a* axis stratifies pro-senescent and profibrotic cell populations in aged kidneys. (A–B). Scatter plots of the linear relationship between log-transformed TPM expression of *Bmal1* (x-axis) and *Cdkn1a* (y-axis) across Zeitgeber Time (ZT) in bulk RNA-Seq samples from young females (A, light orange), old females (A, dark orange), young males (B, light blue), and old males (B, dark blue). Regression lines with 95% confidence intervals are shown, and corresponding Pearson correlation coefficients, adjusted p-values for each cohort is highlighted in boxes with corresponding colors. (C-D). Single-nucleus RNA-Seq data from male (C) and female (D) kidneys at 6 and 24 months, with cells stratified into three populations based on *Bmal1* and *Cdkn1a* expression; ellipses indicate population clusters 1 (green), 2 (red), and 3 (orange). (E-F) KEGG pathway enrichment analysis comparing populations: (E) Top 20 upregulated pathways in population 2 vs. population 1 ranked by significance (-logFDR) with pathways known to be upregulated or altered in senescence underlined in red. (F) Top 20 upregulated pathways in population 3 vs. population 1 ranked by significance (-logFDR) with pathways known to be upregulated or induced by fibrosis underlined in orange.

To investigate this further, we used male and female C57BL/6J mice at 6 and 24 months of age, harvested their kidneys at ZT3, and performed snRNA-seq. After alignment and quality control, we plotted the expression of *Cdkn1a* and *Bmal1* for all nuclei sampled across ages and sexes and identified three distinct populations: cells with a high correlation between the two genes (population 1, green), cells with high *Cdkn1a* expression and low *Bmal1* expression (population 2, red), and cells with high *Bmal1* expression and low *Cdkn1a* expression (population 3, orange) (Figures 5C and 5D). To compare these populations, we performed differential expression analysis using all datasets (6 and 24 month, male and female) and compared population 1 to populations 2 and 3 using Wilcoxon rank-sum tests, with significance defined as adjusted p-value < 0.05. Significant differentially expressed genes (DEGs) were further classified as upregulated (average log2FC ≥ 0.3) or downregulated (average log2FC ≤ –0.3) for KEGG pathway over-representation analysis.

Relative to population 1, population 2 exhibited 142 upregulated and 149 downregulated genes. KEGG enrichment of upregulated genes revealed increased representation of pathways altered or increased during senescence, including Rap1, Ras, MAPK, PI3K–Akt, IgSF CAM, and HIF-1 signaling (Alique et al., 2020; Kumari & Jat, 2021; Lototska et al., 2020) (Figure 5E, red underlined). Downregulated genes were enriched for pathways known to be inhibitory to senescence or downregulated in aging, such as glucagon signaling, taurine metabolism, glyoxylate and dicarboxylate metabolism, renin secretion, and peroxisomes (Cosarderelioglu & Abadir, 2025; Frasca, Saada, Garcia, & Friguet, 2021; Giordano & Terlecky, 2012) (Figure S5A, dark red underlined). Additionally, within the top 10 most significant upregulated genes—*Cdkn1a*, *Trp53cor1*, *Btg2, Ephx1, Ccng1,* and *Creb5*—implicate cell-cycle arrest (*Cdkn1a, Trp53cor1, Btg2*) (J. Wang et al., 2021; Winkler et al., 2022), metabolic dysfunction (*Ephx1*) (Gautheron et al., 2021), and cellular stress (*Ccng1, Creb5*) (A. Wang et al., 2025; Zhang et al., 2024). Together, these findings indicate that population 2 is stressed and cell cycle arrested, and therefore more senescent-like than population 1.

Relative to population 1, population 3 exhibited 281 upregulated and 270 downregulated genes. KEGG enrichment of upregulated genes revealed increased representation of pathways upregulated or involved in fibrosis such as integrin signaling, focal adhesion, cytoskeleton in muscle cells, platelet activation, ECM-receptor interaction, hypertrophic cardiomyopathy, dilated cardiomyopathy, protein digestion and absorption, PI3K-Akt signaling, arrhythmogenic right ventricular cardiomyopathy, and regulation of actin cytoskeleton (Eijgenraam, Sillje, & de Boer, 2020; Rieder et al., 2025) (Figure 5F, orange underlined). Downregulated genes were enriched for pathways typically downregulated in fibrotic cells such as peroxisomes, folate transport and metabolism, glutathione metabolism, pentose and glucuronate interconversions, glyoxylate metabolism, proximal tubule bicarbonate reclamation, PPAR signaling, and pyruvate metabolism (Feng et al., 2023; Rieder et al., 2025) (Figure S5B, dark orange underlined). Notably upregulated genes—*Cfh, Robo2, Lama2, and Rbms3*—indicate enhanced ECM remodeling and a more profibrotic state in population 3 relative to population 1 (Accorsi, Cramer, & Girgenrath, 2020; Fritz & Stefanovic, 2007; Ma, Hua, & Lu, 2025; Zeng et al., 2018). Supporting this interpretation, we also observe significant downregulation of *Ass1*, reduced expression of which promotes fibroblast activation and pulmonary fibrosis, further reinforcing the profibrotic phenotype of population 3 (Li et al., 2021).

We annotated the nuclei for cell type, and found these populations were not a cell type specific phenomenon with nearly all cell types falling into all three populations (Figure S6). Meaning, this strategy of using gene expression uncoupling as a stratification measurement to identify senescent-like and profibrotic states could potentially be cell type universal. This is of interest because cell cycle arrest markers such as *Cdkn1a* have long been used as senescence indicators, yet accumulating evidence shows that their expression alone does not reliably distinguish a truly senescent cell (Suryadevara et al., 2024). Our approach helps pinpoint where that distinction may lie—specifically, in the uncoupling of *Cdkn1a* expression from its circadian counterpart *Bmal1*.

Despite that nuclei were pooled across ages and sexes, pathway-level signatures remained robust even under more stringent significance criteria using an adjusted p-value cutoff of 0.05 rather than 0.1. Critically, classifying nuclei by their *Cdkn1a–Bmal1* expression relationship exposed senescent-like and profibrotic states that marker-only approaches may miss. Together with the bulk RNA-seq observation that *Bmal1–Cdkn1a* expression correlation weakens with age, these results establish a practical circadian–senescence axis for identifying senescent cell states in snRNA-seq data and advance the central paradigm of this work: relational metrics across circadian and senescence genes reveal aging phenotypes even when sampling resolution or effect sizes are limited.

### 2.6 Cultured tail-tip fibroblasts model aging signatures seen *in vivo*

To determine whether age-related cellular changes observed *in vivo* can be modeled in primary cells collected non-invasively, we cultured tail-tip fibroblasts from a subset of the UM-HET3 study animals. For each age and sex, four biological replicates were grown to confluency, synchronized via serum shock, and sampled at seven timepoints over 24 hours starting 16 hours post synchronization (PS) through PS40. Bulk RNA sequencing was performed, reads were aligned using the GBRS pipeline, batch effects were corrected with Learning and Imputation for Mass Spec Bias Reduction (LIMBR), and circadian gene expression was analyzed using the Extended Circadian Harmonic Oscillator (ECHO) application with stringent cutoffs (Period: 20–28 h; Amplitude Change Coefficient > |0.15|; p-value < 0.05; BH/BY FDR < 0.01) to account for genetic diversity (Tables S5-S8). Cohorts were analyzed by age and sex and then compared. Consistent with prior *in vivo* reports in male C57BL/6J mice, fibroblasts from young animals of both sexes exhibited a greater number of circadian genes than fibroblasts from aged animals, and each age group displayed unique circadian signatures, indicating that circadian regulation shifts across the lifespan (Figures 5A, 5B, S7) (Wolff et al., 2023). To identify pathways undergoing age-dependent circadian regulation, we examined biological process ontologies associated with circadian genes in young and old fibroblasts. Ontologies were reduced using rrvigo with a threshold of 0.7 and ranked by adjusted p-value with the top 5 pathways selected for visualization.

In fibroblasts from young males, circadian genes were enriched for DNA replication, ribonucleoprotein complex biogenesis, regulation of cell cycle phase transition, and chromosome segregation—processes essential for proliferation and repair (Figure 6C). While core pathways such as DNA replication and ribonucleoprotein biogenesis remained circadian in aged males, additional enrichment emerged for ribosome biogenesis and protein localization (Figure 6D). In fibroblasts from young females, circadian genes were enriched for macroautophagy, cellular component disassembly, cell cycle, and ribosome biogenesis, suggesting robust transcriptional control and organelle maintenance (Figure 6E). In aged females, many pathways persisted, but enrichment shifted toward regulation of cell cycle phase transition and cellular component disassembly (Figure 6F). Notably, ribonucleoprotein complex biogenesis, macroautophagy, and cell cycle processes were enriched in circadian genes in fibroblasts and previously emerged as the top variably expressed pathway in aged kidneys (Figures 6C-6F, S4).

**Figure 6.**
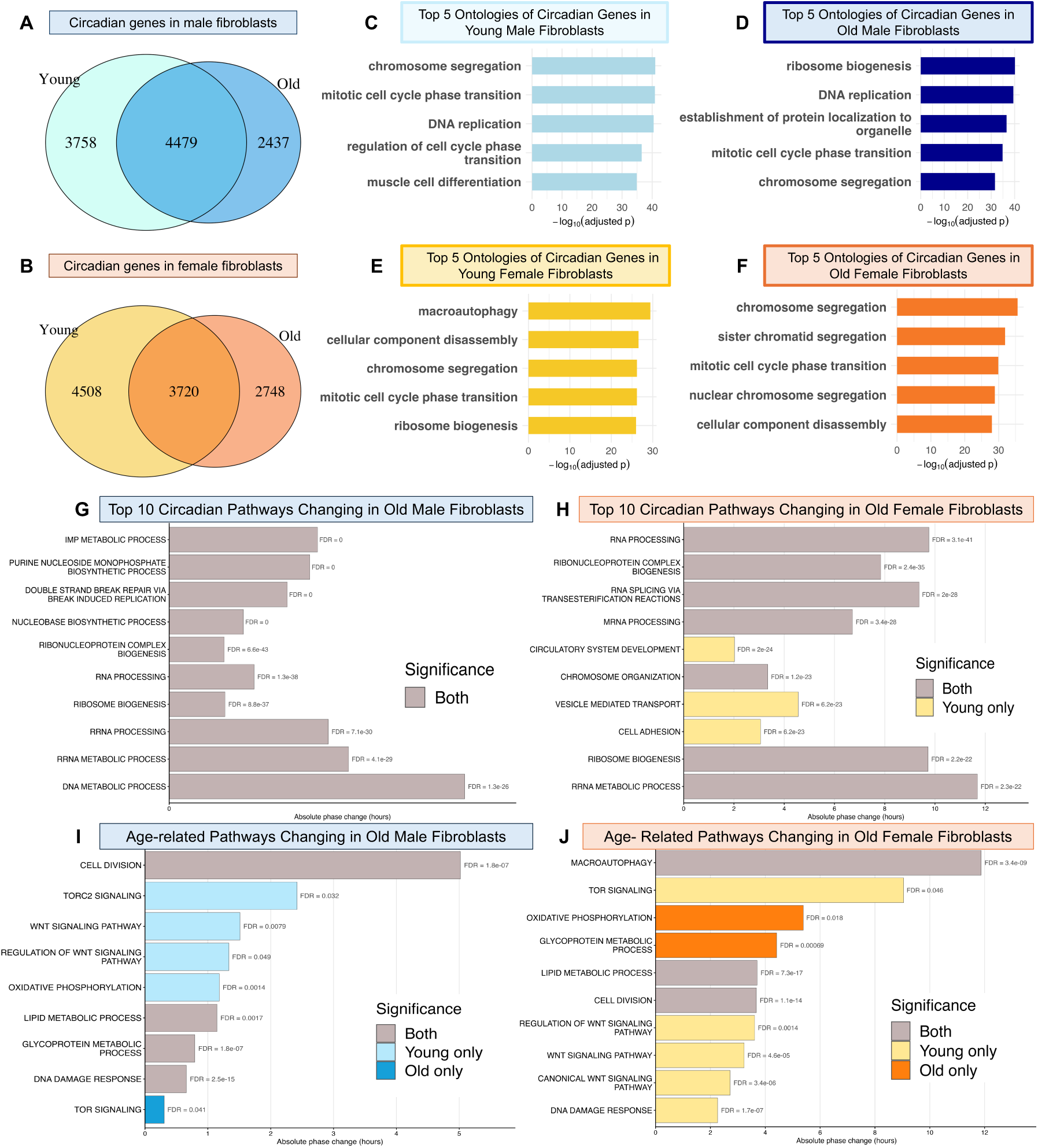
Age-related alterations to circadian pathways are recapitulated in fibroblast cell culture. (A & B) Venn diagrams depicting the number of circadian genes found in young and old male fibroblasts (A) and young and old female fibroblasts (B). (C-F). Top 5 enriched Gene Ontology biological processes by -log(adjusted p-value) among circadian genes in young male (C), old male (D), young females (E), and old female (F) fibroblasts. (G–J) Bar graphs summarize pathway-level changes in circadian phase with aging. Panels G and H show pathways that are circadianly expressed at an enriched phase in young (light colors), old (dark colors), or both (gray), ranked by most significant false discovery rate (FDR) values for males (G) and females (H). Panels K and L highlight selected aging-specific pathways that were significantly enriched in aged fibroblasts for males (K) and females (L). Bar lengths represent magnitude of phase change. FDR for each pathway is reported to the right of the bar.

Ontology analysis of circadian genes unique to young or old samples further revealed that young-specific circadian pathways aligned with tissue-specialized functions (Figure S7, light blue and yellow), whereas aging-specific circadian pathways centered on ribosome biogenesis, macroautophagy, and cell-cycle regulation (Figure S7, dark blue and orange)—mirroring pathways showing increased variability in aged kidneys (Figure S4). This pattern suggests a loss of tissue-specific circadian entrainment with aging and a shift toward repair-associated programs, consistent with prior profiling of age-related circadian changes (Wolff et al., 2023). These findings indicate that fibroblast cultures retain tissue specific signatures in culture and can model aspects of *in vivo* aging biology.

To assess temporal changes to pathways in aged fibroblasts, we performed PSEA on circadian genes and compared significant pathways between young and old samples. In males, the most significantly phase changing pathways in aging were inosine monophosphate (IMP) metabolism, nucleobase synthesis, and DNA repair, though phase shifts were modest (<1hr) (Figure 6G). The largest shifts occurred in hormone stimulus, immune response, and fatty acid catabolism pathways (Figure S8, blue). In females, significantly changing pathways in aging involved RNA processing, splicing, and chromosome organization, with phase shifts ranging from 2-11hrs (Figure 6H). The largest phase shifts were in actin filament organization, potassium ion transport, fat cell differentiation, and macroautophagy (Figure S8, orange). Together, these results show that aging alters circadian control of pathways linked to cellular stress responses and homeostasis in both sexes, but diverges, with males showing more metabolic, immune, and DNA repair changes, while females exhibit shifts in RNA processing, cytoskeletal organization, and autophagy—revealing sex-specific circadian aging trajectories.

Moreover, curated kidney aging pathways, including cell division, TOR signaling, Wnt signaling, oxidative phosphorylation, lipid metabolism, glycoprotein metabolism, and DNA damage response, also showed significant phase shifts in aged, cultured fibroblasts (Figures 6I and 6J). These parallels indicate that renal age-related circadian dysregulation can be partially recapitulated in primary cell culture, with both sexes showing coordinated changes in cell-cycle regulation, mTOR signaling, and energy metabolism both *in vivo* and *in vitro* (Figures 4A, 4B, S4). However, the tissue-specific nature of the most enriched pathways indicates that tail-tip fibroblasts maintain their intrinsic identity, enabling them to capture general aging signatures yet constraining their ability to model tissue-unique aging alterations.

## 3. Discussion

A central finding of this study is that aging alters the architecture of the renal circadian clock, revealing that the most consequential circadian changes in aging are not simply dampened rhythms but the erosion of coordinated timing within the oscillator itself. Rather than losing rhythmicity outright, aged kidneys show a breakdown in the canonical anti-phase relationship between core clock components, a form of circadian dysregulation that exposes vulnerabilities within the timing network that are not detectable through amplitude-based assessments alone. This disruption of internal timing provides a potential mechanistic explanation for why aging tissues can appear rhythmically intact when individual genes are analyzed in isolation yet functionally compromised when their interactions are considered.

Building on this insight, our work introduces the concept of circadian uncoupling—the loss of coordinated timing between the clock and its downstream targets—as a powerful, relational framework for identifying altered cellular states. We demonstrate this using the *Bmal1–Cdkn1a* expression axis, showing that aging weakens the temporal partnership between a core clock activator and a key senescence effector. This uncoupling stratifies distinct senescent-like and pro-fibrotic populations at the single-cell level. Importantly, this relational decay emerges even when both genes remain expressed and oscillatory, highlighting that the breakdown of timing relationships can be more informative than changes in absolute expression levels. As such, the uncoupling metric provides a higher-order view of cellular state that traditional markers cannot capture.

Although we apply this framework to a single axis in kidney aging, the broader conceptual potential of circadian uncoupling is substantial. Core clock components regulate thousands of rhythmic outputs across nearly every tissue, including pathways involved in metabolism, immune signaling, and mitochondrial function (García Cobarro et al., 2025). Because circadian disruption contributes to multiple age-associated conditions, including cancer, neurodegenerative disease, and cardiovascular dysfunction, this relational approach could be extended to other clock–target gene pairs to reveal early or transitional cellular states relevant to these contexts (García Cobarro et al., 2025). In this way, circadian uncoupling provides a generalizable strategy for detecting altered cell states that may be invisible using standard markers or single-timepoint profiling.

By resolving circadian and sex effects in young and old kidneys, we defined how senescence phenotypes vary across the day and how these patterns align with broader pathway-level changes. Males showed daytime inflammatory SASP peaks (ZT3, ZT9), which corresponded to increased immune ontologies at the same timepoints, followed by a nighttime shift toward metabolic pathways (ZT15, ZT21). In contrast, females displayed sustained inflammatory and ECM-remodeling signatures across the day, with a pronounced nighttime cytokine/chemokine SASP at ZT15—mirrored by persistent immune ontologies at all timepoints. This tight concordance between SASP timing and immune pathway enrichment suggests that inconsistencies in senescence marker detection across studies may stem from their circadian regulation. Demonstrating these sex- and time-structured signatures in a genetically diverse model system underscores circadian timing as a critical dimension for interpreting kidney aging phenotypes.

Our analysis of age-related variability revealed widespread instability in pathways central to cellular homeostasis, including translation, RNA processing, energy metabolism, macroautophagy, and TOR signaling, consistent with hallmark aging processes (Lopez-Otin et al., 2023). Similarly, pathways that were more variably expressed *in vivo* showed age-related phase shifts *in vitro*: synchronized old tail-tip fibroblasts exhibited reduced circadian regulation and age-dependent shifts in key pathways such as mTOR signaling, cell-cycle, and oxidative phosphorylation. Although these changes were present in both sexes, females showed larger phase shifts in aged fibroblasts and a greater number of variably expressed genes in aged kidney tissue compared to males. While cultured fibroblasts effectively model major age- and sex-specific circadian changes, they also highlight that tissue-specific oscillatory programs diverge with age. Together, these findings indicate that aging increases transcriptional noise and reorganization of circadian-regulated pathways across tissues, while cultured fibroblasts offer a practical but inherently partial window into how circadian timing deteriorates during aging.

This study has several notable strengths, including multi-Zeitgeber sampling in kidney tissue, sex-stratified analyses, explicit modeling of variance with missMethyl, validation of relational metrics across bulk and single-nucleus datasets, and paired time-course profiling in synchronized primary fibroblasts. However, several limitations remain: temporal resolution in kidney samples was restricted to four ZTs, replicate numbers were sex-imbalanced at certain aged timepoints, and single-nucleus data were collected at only one circadian time point. These constraints were mitigated by using relational metrics to capture circadian dysregulation (*Bmal1–Per2*) and uncoupling (*Bmal1–Cdkn1a*), both of which remain informative even with limited temporal sampling and provide a practical framework for studies constrained to one or a few timepoints.

Together, these findings show that aging degrades circadian regulation, producing sex- and time-structured senescence-associated gene expression and increasing variability in clock-regulated pathways. By applying a clock-informed relational metric, the *Bmal1–Cdkn1a* axis, we identified senescent-like and profibrotic states in snRNA-seq data that are undetectable using single marker detection, while primary fibroblasts reproduced age-dependent phase shifts in nutrient and mitochondrial pathways. These results highlight circadian timing as a fundamental dimension of aging biology and demonstrate that relational metrics such as circadian uncoupling provide a scalable and practical framework for detecting altered cellular states.

## 4. Methods

All software tools, pipelines, and statistical frameworks used in this study are fully cited in the Supporting Methods to maintain clarity and readability of the main text.

### 4.1 Animals

UM-HET3 animals (strain #036603) were obtained from The Jackson Laboratory (JAX). A cohort of 30 males and 31 females was aged to 24 months. Additionally, 32 males were acquired and aged to 29 weeks, and 32 females were obtained and aged to 27 weeks. C57BL/6J animals (strain #000664) were obtained from JAX at 6 and 24 months of age (n=3 per sex and age). Throughout the study, all animals were housed in the same room, provided with standard 6% fat chow (5KOG Lab Diet, St. Louis, MO) and water ad libitum, given nestlet enrichment, and maintained on a 12-hour light/12-hour dark cycle (lights on at 6:00 AM, off at 6:00 PM). All procedures were approved by the JAX Institutional Animal Care and Use Committee (protocol #22-141).

### 4.2 Circadian tissue collection

Animals were harvested every 6 hours across a 24-hour cycle, beginning at 9:00 AM. Lights in the housing room turned on at 6:00 AM (ZT0), yielding sampling timepoints at ZT3, ZT9, ZT15, and ZT21. Animals were euthanized by cervical dislocation, and tissues were immediately collected and flash-frozen. For nighttime collections, mice were transferred to a dark procedure room (lights off at 6:00 PM) with red-filtered coverings to prevent light exposure, and tissues were harvested under white lamp illumination.

### 4.3 Cell culturing and circadian synchronization

Tail-tip fibroblasts were derived from 8 animals per harvest (4 young, 4 old) at ZT3 for both sexes. Tails were put into 15ml conicals with 13ml of DMEM media, 100X Penstrep (1:100) and Beta-mercaptoethanol (BME, 1:1000), then minced and divided into six centrifuge tubes containing 400μl of the same media. Collagenase D solution (400μl; 5mg/ml) was added, and tubes were shaken at maximum speed for 110 min at room temperature. Tail pieces were transferred to 15ml conicals containing 4mls of growth media containing DMEM, fetal bovine serum (FBS, 1:10), 100X Penstrep (1:100), and BME (1:1000), further disassociated with a P1000 pipette, allowed to settle, and 2ml of media was removed to eliminate remaining hair. An additional 2ml of growth media was added, tissue was pipetted with a P200 pipette, then centrifuged at 1400rpm for 5 min at room temperature. Supernatant was removed, tissue was resuspended in 12ml of growth media, plated in 6-well plates, and cultured at 37°C, 3% O_2_ with media changes imaging every 3 days.

Circadian synchronization followed established protocols (Collins et al., 2021). Young fibroblasts were synchronized after 9 days in culture and old fibroblasts after 10 days, once confluent. Cell received complete media with low-glucose DMEM (1g/L), FBS (1:10), 100X Penstrep (1:100), and BME (1:1000) for 24hrs, followed by 24hrs in starvation media (low glucose DMEM, 100X Penstrep 1:100, and BME 1:1000). Cells were then exposed to shock media (low glucose DMEM, FBS 1:2, 100X Penstrep 1:100, and BME 1:1000) for 2hrs, washed with warm PBS, and returned to complete media. Sixteen hours later, fibroblasts were harvested every 4hrs for 24hrs. At each timepoint. cells were washed with cold PBS, scraped, pelleted at 500 *x g* for 5min at room temperature, and flash frozen.

### 4.4 Bulk RNA-sequencing of whole-kidneys and tail-tip fibroblasts

Frozen tissues were ground in liquid nitrogen with a mortar and pestle. Cell pellets were harvested and flash frozen with three technical replicates per timepoint. Aliquots of ground tissue and three cell pellets per animal per timepoint were sent to Novogene (Sacremento, CA, USA) for RNA extraction with poly A enrichment. Libraries were sequenced on a NovaSeq X Plus (150bp read length, 30M depth). Raw FASTQ files were aligned with the GBRS pipeline (https://github.com/churchill-lab/gbrs) to generate raw counts. These counts were used for downstream analysis; however, young male kidney samples were batch corrected using ComBat due to multiple sequencing runs ahead of all downstream analysis.

### 4.5 Cosinor analysis of whole kidneys

To identify circadian oscillations in the kidney, we used GLMMcosinor analysis with DHARMa residual diagnostics. A gene was classified as circadian if it had significant sine or cosine, amplitude, and phase estimates, and if DHARMa residuals showed no significant deviation (quantile p-value > 0.05), indicating an adequate model fit.

### 4.6 Differential expression analysis of whole kidneys

Raw counts were analyzed in DESeq2 using a minimum threshold of 10 counts in at least 5 samples, with batch, age, and timepoint included as variables. Differential expression as tested using Wald statistics, comparing 6 vs 24 months samples within each sex and timepoint.

### 4.7 Ontology and pathway analysis

For the whole kidney datasets, DEGs between young and old UM-HET3 males and females were included in enrichment analysis if they met the following criteria: upregulated genes with a log₂FC> 1 and downregulated genes with a log₂FC< –0.5, each with an adjusted p-value ≤ 0.1. For the tail-tip fibroblast datasets, genes were included in the enrichment analysis they were determined to be circadian as described above. Enrichment for both the kidney and fibroblast datasets were run using clusterProfiler with a q value < 0.05 for Biological Process and Molecular Function ontologies. Resulting terms were clustered with rrvgo using “Rel” semantic similarity and a 0.5 similarity threshold. The top five Biological Process ontologies were selected based on the most significant false discovery rate (FDR) values.

### 4.8 Variance analysis

Variance analysis was performed using the missMethyl package (v1.42.0), which contains functions to measure variability in RNA sequencing data as previously described in Wolff et al. 2023. Raw RNA-seq counts were analyzed using edgeR (v4.6.3) to create a DEG list and normalized with voom (quantile normalization) in limma (v3.64.3) to model the mean–variance relationship and compute the precision weights for linear modeling. For each analysis, linear models were fitted with age as the primary factor and timepoint as a random effect using duplicate correlation in limma to account for repeated circadian measurements. For merged analyses of both sexes, sex was included as a fixed effect.

DV was assessed using missMethyl’s varFit and topVar functions which apply Levene’s test-based models to detect variance differences between age groups. Genes with DV FDR < 0.05 were considered significant. Analyses were run separately for males and females, as well as combined, and variance contributions from sex, age, timepoint, and batch were examined to confirm that age and sex were the dominant sources of variability.

#### 4.8.1 Pathway enrichment

DV-associated genes were analyzed using clusterProfiler (v4.16.0) with Benjamini–Hochberg (BH) correction. Redundant GO terms were collapsed with rrvgo (v1.10.0) using “Rel” semantic similarity and a 0.9 similarity threshold, retaining terms with dispensability ≤ 0.1 with the org.Mm.eg.db annotation database (v3.21.0). The top 20 pathways based on most significant -log₁₀ adjusted p-value and were visualized with ggplot2 (v3.5.2).

### 4.9 snRNA-Seq of kidneys

Kidneys from 6 and 24month male and female C57BL/6J mice were harvested at ZT3 following a published protocol (Robinson, Sheehan, Garland, & Korstanje). Nuclei were isolated using establish methods (L Daigle, A Perry, F Flynn, & T Courtois, 2024), then encapsulated and barcoded using the 10x Genomics Chromium Single Cell 3’ platform following manufacturer protocols and prior published work (L. Daigle et al., 2024). Libraries were sequenced on an Illumina NovaSeq X + 10B (100-cycle flow cell). Illumina base call (BCL) files were converted to FASTQ format using ‘bcl2fastq’ v2.20.0.422 (Illumina).

FASTQ files were aligned to GRCm38 with GENCODE vM23 (10x Genomics mm10 reference 2020-A) using the 10x Genomics Cell Ranger pipeline. Quality control (QC) removed low-quality nuclei and technical artifacts (< 200 detected genes, >10% mitochondrial reads, or >25% ribosomal reads). Genes detected in <3 nuclei were excluded. Nuclei with <500 unique molecular identifiers (UMIs), >25,000 UMIs, or >6,000 detected genes were removed to limit doublets and low-quality profiles. Additional doublet detection was performed using DoubletFinder (v2.0.3).

After QC, 118,860 high-quality nuclei remained. Preprocessing and analysis were performed in R using Seurat (v4.0.1), including log-normalization, principal component analysis (PCA), uniform manifold approximation and projection (UMAP), and unsupervised clustering. Cluster marker genes were identified using the Wilcoxon rank-sum test, and cell types were annotated using a mouse kidney reference (Clark et al., 2019).

### 4.10 Identification of *Bmal1*- and *Cdkn1a-*expressing cell populations

To characterize heterogeneous expression patterns of *Bmal1* (*Arntl*) and *Cdkn1a*, we quantified co-expression across all nuclei and stratified them into three populations. Population 1 contained nuclei with the strongest *Bmal1* and *Cdkn1a* expression correlation. Population 2 included cells with disproportionately higher *Cdkn1a* relative to *Bmal1* (y > x + ε) and Population 3 included cells with disproportionately higher *Bmal1* relative to *Cdkn1a* (y < x – ε), where ε defines the bandwidth around the Population 1 diagonal representing the strongest co-expression.

DEGs between Population 1 and Populations 2–3 were identified in Seurat using the Wilcoxon rank-sum test (adjusted p-value < 0.05). Significant DEGs were stratified into upregulated (avg_log2FC ≥ 0.3) and downregulated (avg_log2FC ≤ –0.3) gene sets for pathway analysis. KEGG pathway over-representation analysis (ORA) was performed in R (v 4.5.0) using the clusterProfiler with org.Mm.eg.db for mouse annotation. Gene symbols were converted to Entrez IDs, and ORA was run separately on up- and downregulated sets. Multiple-testing correction used the BH method, with enriched pathways defined as p-value < 0.05 and q-value < 0.05.

### 4.11 Circadian gene expression analysis of tail tip fibroblasts

Cells were processed, sequenced, and aligned as described above. Raw counts were converted to TPM, and batch corrected per group (young males, old males, young females, old females), with all timepoints and replicates processed together using LIMBR. Circadian analysis was performed with ECHO v4 in free run mode, using 12 paired replicates per timepoint, with smoothing, removal of unexpressed genes, and linear detrending. Genes were classified as circadian if they had a period of 20-28hrs, AC coefficient >0.15 or <-0.15, p value <0.05, BH and Benjamini–Yekutieli (BY) p-value < 0.01.

### 4.12 PSEA

Circadian genes were obtained from prior ECHO analysis. Circadian phase was derived from each gene’s Hours Shifted scaled by its estimated period. Phases were standardized to a 24hrs, and initial amplitude was used as a nonnegative weight when computing pathway-level mean phases. Gene sets were sourced from MSigDB C5 GO Biological Process for *Mus musculus* via the msigdbr package. For each age group, MSigDB gene symbols were intersected with rhythmic genes, retaining sets with ≥10 overlapping genes. For each pathway, gene phases were converted to radians and the Rayleigh test for circular nonuniformity was applied using the circular package. Weighted circular means were computed as:

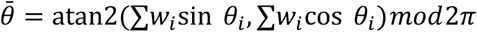

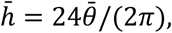

where *w_i_* denotes amplitude weights, *θi* denotes peak phase, and θˉ is the weighted circular mean. Peak phase was converted from radians into circadian time by solving for hˉ. For each pathway, we recorded gene counts, mean peak phase (hrs), Rayleigh *Z*, and p-value, adjusting across pathways using BH FDR; pathways with FDR < 0.05 were considered significantly clustered. Age-related phase shifts were calculated by pairing pathways present in both groups and computing the shortest signed circular difference within [−12, +12]:

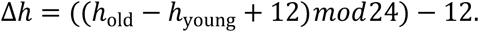

We calculated ∣ Δℎ ∣ for each pathway, retained those with FDR < 0.05 in any group, and categorized significance as “Both”, “Young only”, or “Old only”.

## Author contributions

G.T.C. and R.K. contributed to conceptualization and methodology, leading the study design and experimental planning. R.K. and N.A.R. were responsible for funding acquisition and supervision, providing oversight for data interpretation. G.T.C. prepared the writing of the original draft, and R.K. contributed to writing and review and editing through critical revision. G.T.C. and R.K., together with C.M.F., R.E.R., C.W., S.S., A.W., S.S., and A.B., contributed to investigation, including experimental work, data acquisition, and manuscript editing. G.T.C. and Y.Y. performed formal analysis and contributed to data interpretation.

## Acknowledgements

We thank the Single Cell Biology, Genome Technologies, Data Science, and Cyberinfrastructure high-performance computing teams at The Jackson Laboratory for their expert support throughout this work. These shared resources are partially funded by the JAX Cancer Center (P30 CA034196).

## Data availability statement

All data will be submitted and publicly available on the Cellular Senescence Network Consortium (SenNet) data portal at https://data.sennetconsortium.org/.

