## Supplemental Information for "Circadian Dysregulation in Aging Alters Senescence and Inflammatory Pathways in a Sex- and Time-of-Day–Dependent Manner"

### Supporting Methods

To improve readability, all software tools, computational pipelines, normalization methods, and statistical frameworks are provided here rather than in the main text. These tools supported data processing, genome alignment, normalization, circadian modeling, variance analysis, and pathway enrichment.

#### Alignment, normalization, and batch correction

RNA-seq alignment and genotype reconstruction used the GBRS pipeline (Choi et al., 2025). Transcript abundance was normalized to transcripts per million (TPM) (Wagner, Kin, & Lynch, 2012). Batch correction for whole-kidney RNA-seq counts used ComBat-seq (Y. Zhang, Parmigiani, & Johnson, 2020).

#### Circadian modeling

Circadian oscillations were analyzed using GLMMcosinor (Cornelissen, 2014; Parsons, Jayasinghe, White, Chunduri, & Rawashdeh, 2024), and DHARMa residual diagnostics (Hartig, 2024). Fibroblast RNA-seq data underwent batch removal using LIMBR (Crowell, Greene, Loros, & Dunlap, 2019), and circadian oscillations were identified using ECHO (De Los Santos et al., 2020).

#### Differential gene expression of whole-kidney tissue

Differential expression analyses used DESeq2 (Love, Huber, & Anders, 2014).

#### Variance analysis

Count normalization and dispersion modeling included edgeR (Robinson, McCarthy, & Smyth, 2010) and limma/voom (Ritchie et al., 2015). Differential variability was assessed with missMethyl (Phipson, Maksimovic, & Oshlack, 2016).

#### Gene ontology and pathway analysis

For Gene Ontology enrichment we used clusterProfiler (Yu, Wang, Han, & He, 2012). Redundant GO terms were reduced using rrvgo (Sayols, 2023). MSigDB gene sets were accessed via msigdbr (Dolgalev, 2025). For pathway analysis in snRNA-Seq data, we used KEGG (Kanehisa, Furumichi, Sato, Matsuura, & Ishiguro-Watanabe, 2025).

#### Single-nucleus RNA-Seq processing

Kidney snRNA-seq reads were processed with Cell Ranger (Zheng et al., 2017). Doublet detection used DoubletFinder (McGinnis, Murrow, & Gartner, 2019). Downstream filtering, normalization, and clustering were performed using Seurat (Satija, Farrell, Gennert, Schier, & Regev, 2015).

#### Pathway and enrichment analyses

Phase Set Enrichment Analysis on tail-tip fibroblasts followed established methodology (R. Zhang, Podtelezhnikov, Hogenesch, & Anafi, 2016).

**Supporting Figures**

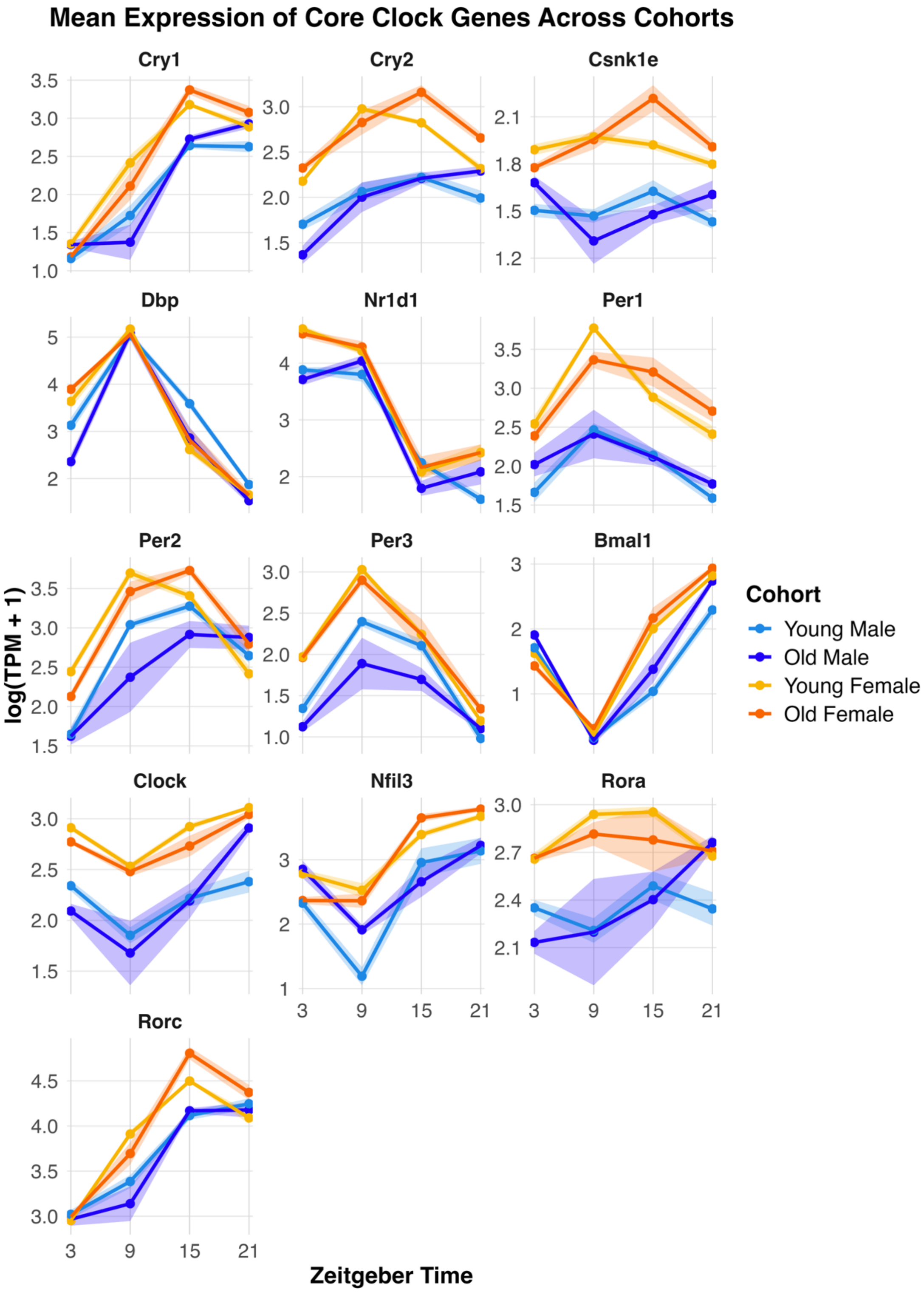


**Figure S1. Core clock gene expression across cohorts.** Core clock genes plotted in mean log(TPM+1) with +-sem shaded over Zeitgeber Time (ZT).

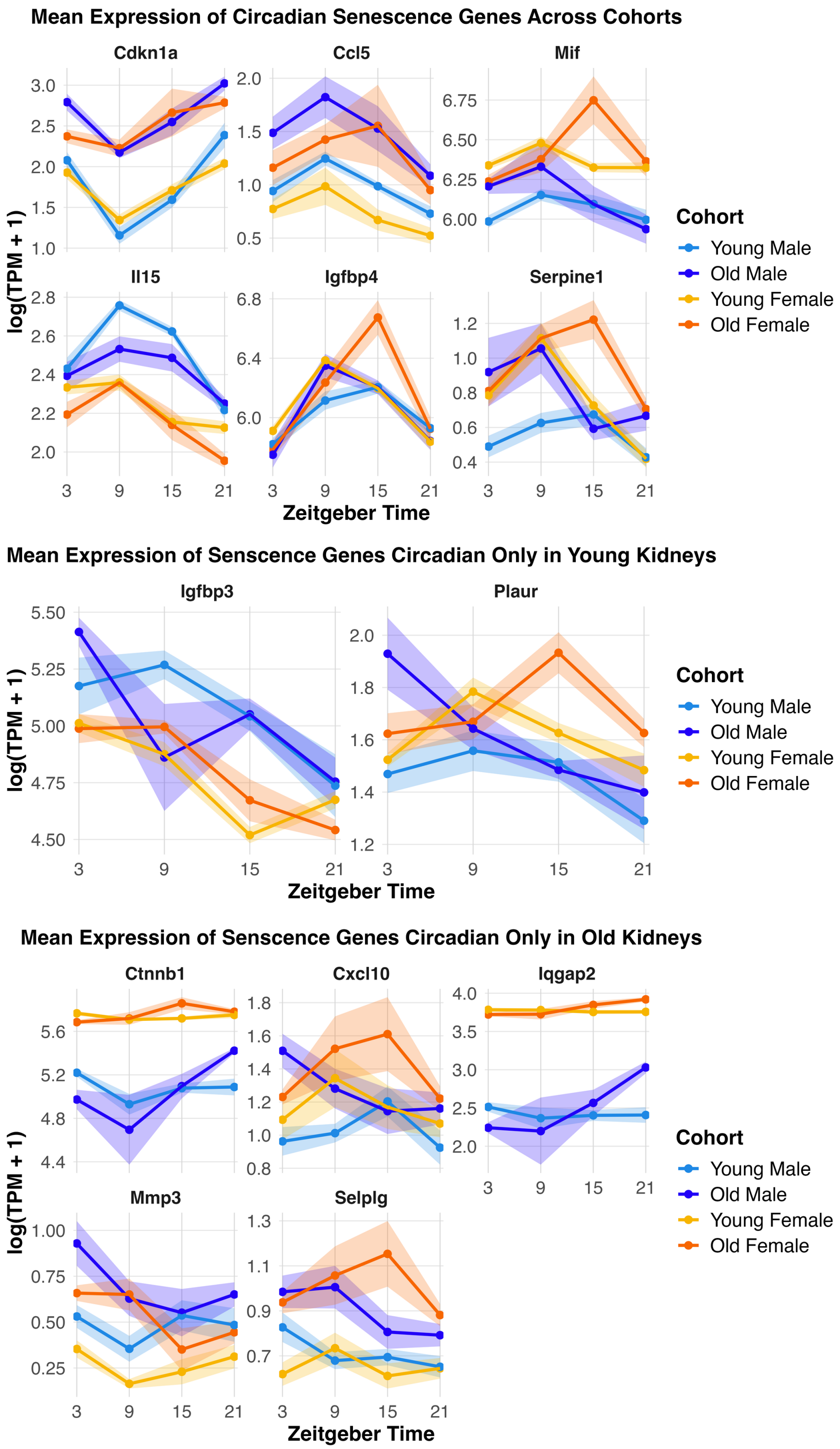


**Figure S2. Circadian senescence gene expression across cohorts.** Senescence genes plotted in mean log(TPM+1) with +-sem shaded over Zeitgeber Time (ZT).

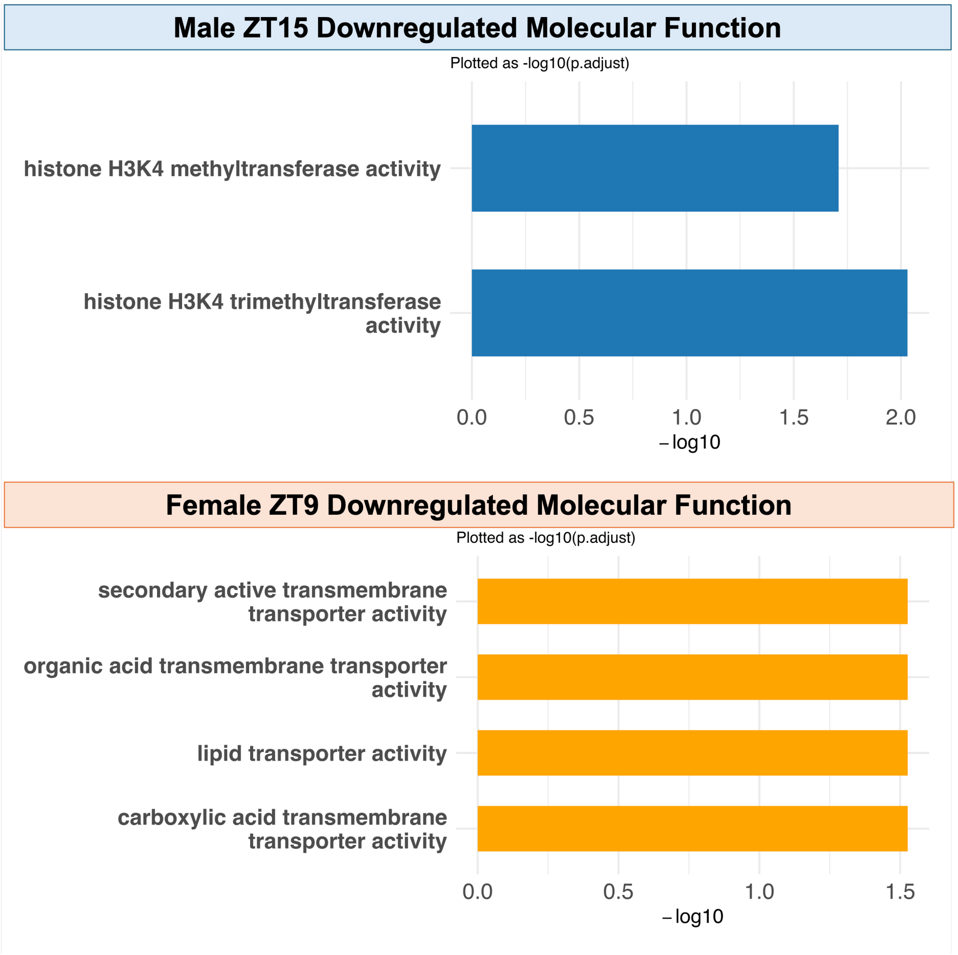


**Figure S3. Downregulated molecular function ontologies.** At ZT15 in males and ZT9 in females, no significantly downregulated Biological Process ontologies were detected. To further investigate this timepoint, we quantified enriched Molecular Function ontologies and plotted the –log10 transformed enrichment significance.

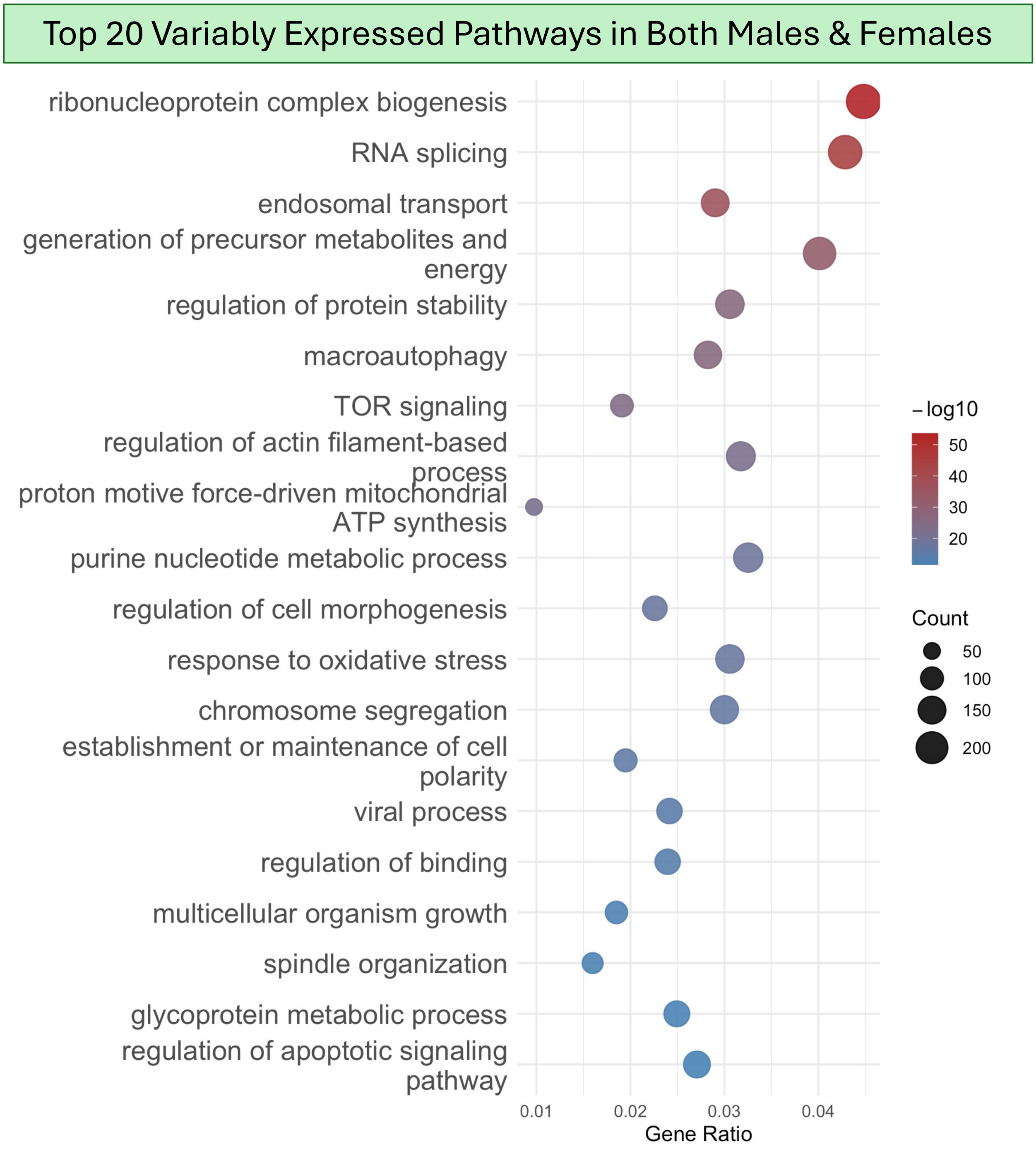


**Figure S4. Top 20 variably expressed genes in both males and females.** Combined male and female data showed the above top 20 pathways being variably expressed in aging. Gene counts are shown in size of dot, and significance is colored with most significant colored as red and least significant colored in blue.

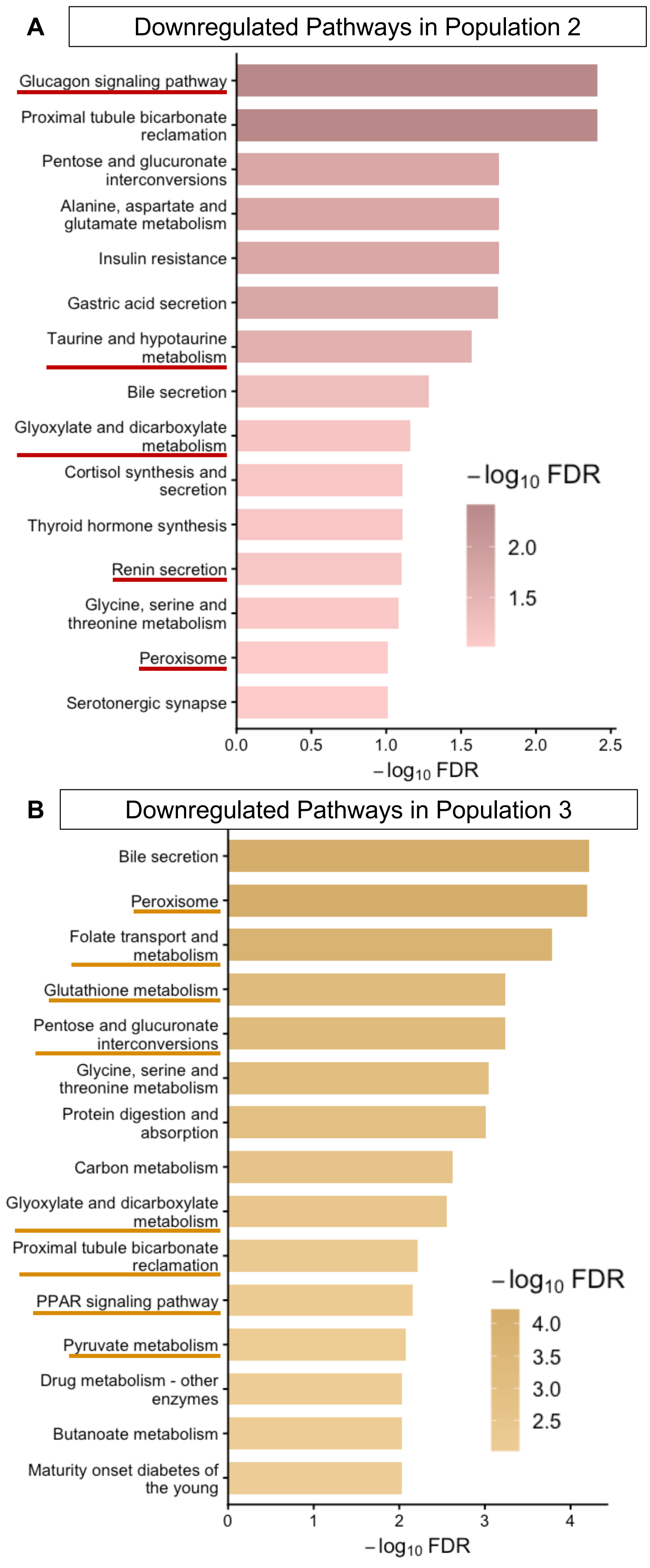

A
Downregulated Pathways in Population 2
Glucagon signaling pathway
Proximal tubule bicarbonate reclamation
Pentose and glucuronate interconversions
Alanine, aspartate and glutamate metabolism
Insulin resistance
Gastric acid secretion
Taurine and hypotaurine metabolism
Bile secretion
Glyoxylate and dicarboxylate metabolism
Cortisol synthesis and secretion
Thyroid hormone synthesis
Renin secretion
Glycine, serine and threonine metabolism
Peroxisome
Serotonergic synapse
−log10 FDR
2.0
1.5
0.0
0.5
1.0
1.5
2.0
2.5
−log10 FDR
B
Downregulated Pathways in Population 3
Bile secretion
Peroxisome
Folate transport and metabolism
Glutathione metabolism
Pentose and glucuronate interconversions
Glycine, serine and threonine metabolism
Protein digestion and absorption
Carbon metabolism
Glyoxylate and dicarboxylate metabolism
Proximal tubule bicarbonate reclamation
PPAR signaling pathway
Pyruvate metabolism
Drug metabolism - other enzymes
Butanoate metabolism
Maturity onset diabetes of the young
−log10 FDR
4.0
3.5
3.0
2.5
0
1
2
3
4
−log10 FDR

**Figure S5. Downregulated pathways in populations 2 and 3 compared to population 1.** A). Top 20 downregulated pathways in population 2 compared to population 1 plotted by most significant FDR with pathways known to be downregulated in senescence underlined in dark red. B). ). Top 20 downregulated pathways in population 3 compared to population 1 plotted by most significant FDR. With pathways known to be downregulated in fibrosis underlined in dark orange.

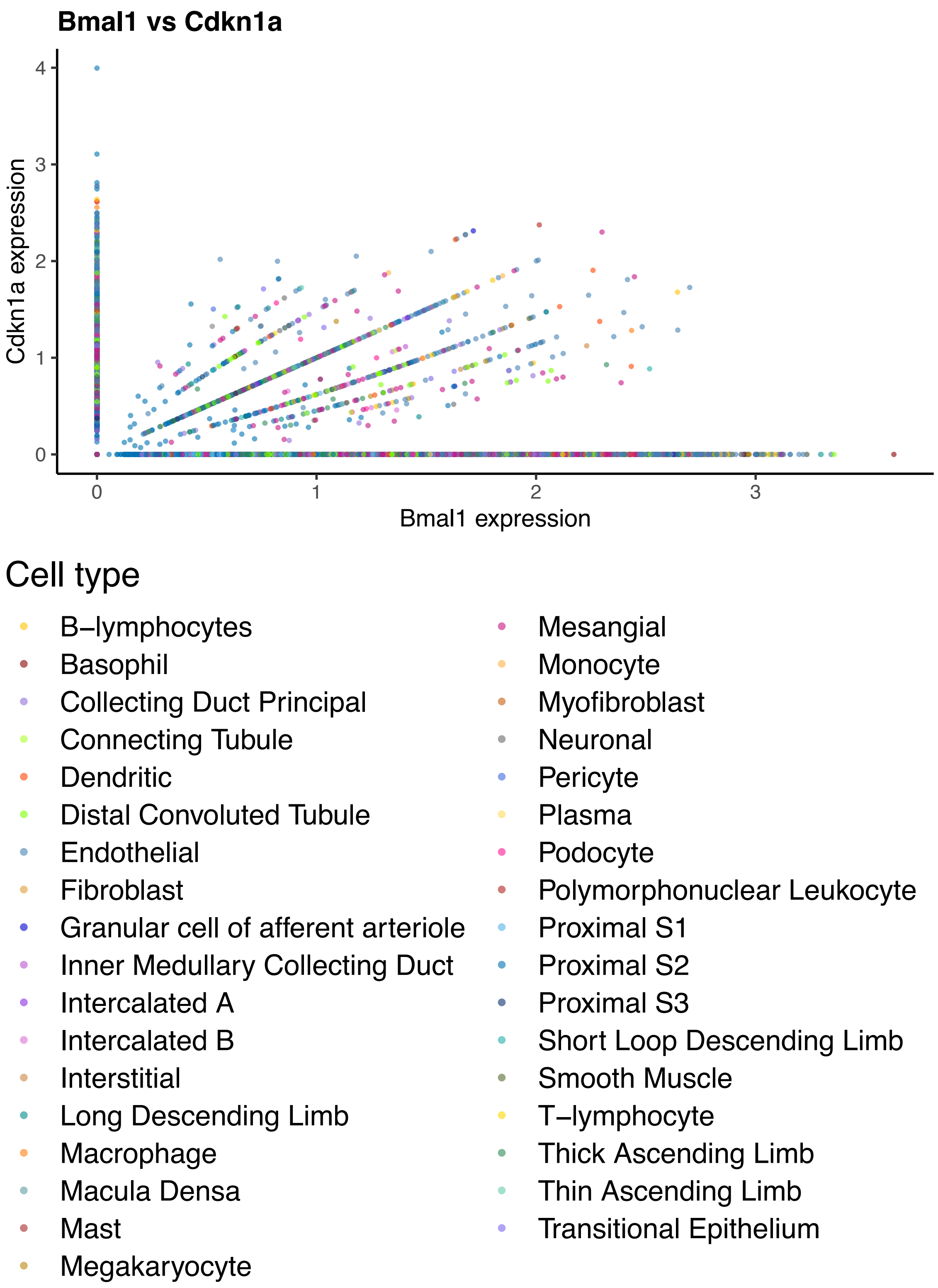


**Figure S6. Cell types of nuclei plotted for snRNA-Seq.** snRNA-Seq log normalized expression of *Bmal1* and *Cdkn1a* with cell type highlighted by color.

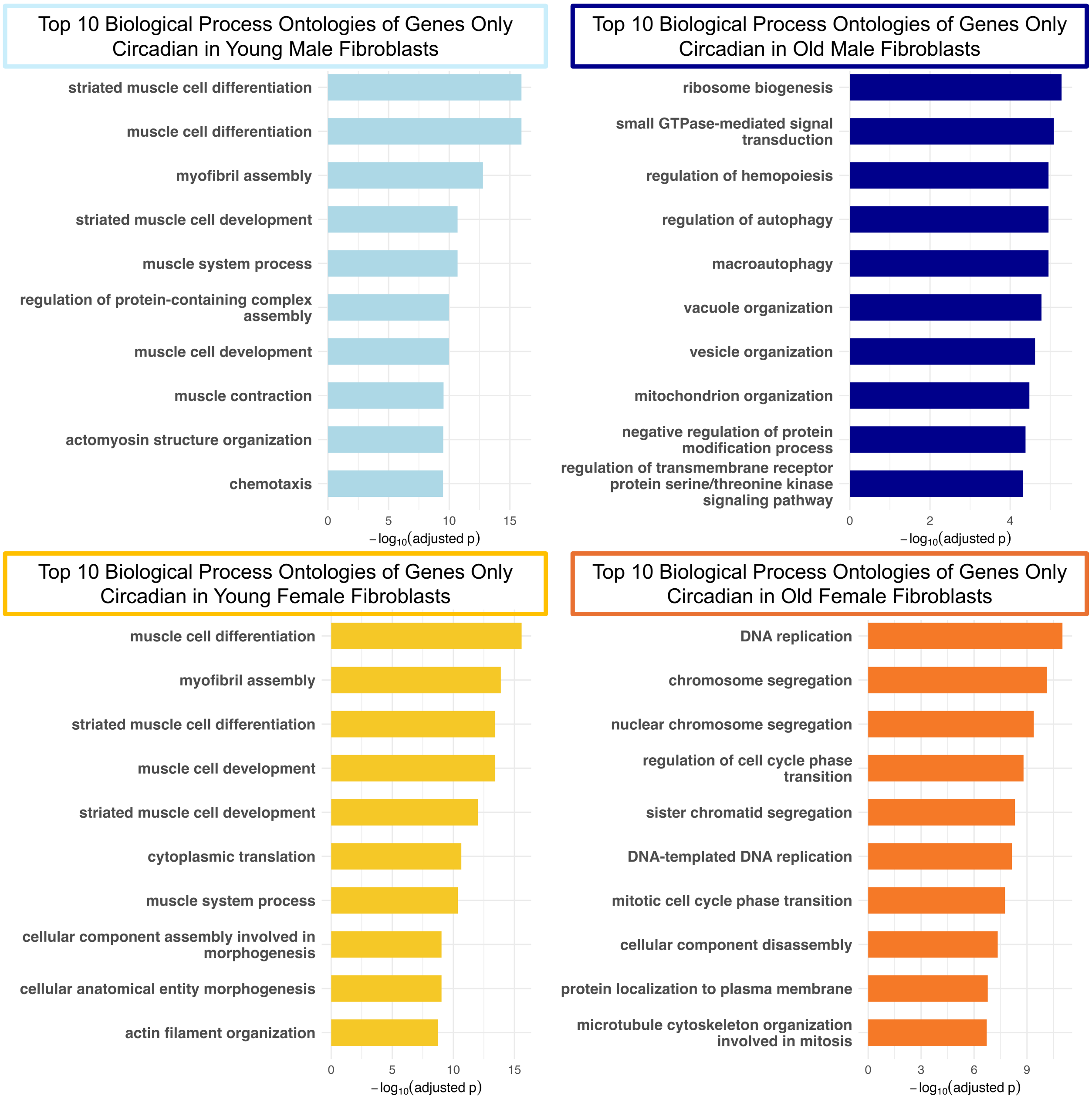


**Figure S7. Ontologies of circadian genes in only in young and old, male and female tail-tip fibroblasts.** Top 10 most significant biological process ontologies by -log(adjusted p value) of genes that are only circadian in young male fibroblasts (light blue), old male fibroblasts (dark blue), young female fibroblasts (yellow), and old female fibroblasts (dark orange).

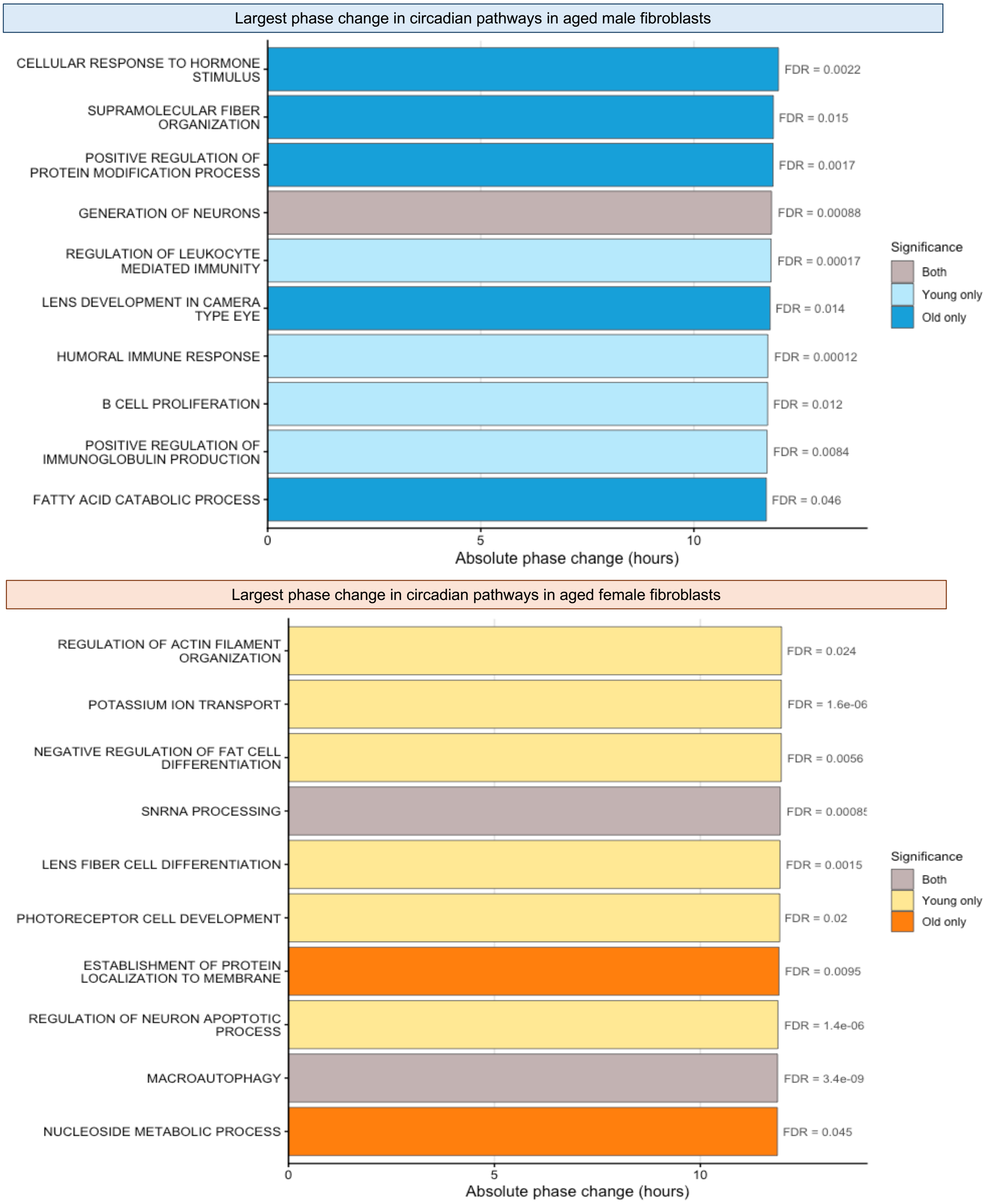


**Figure S8. Top pathways with the largest phase shifts in male and female tail-tip fibroblasts.** The top 10 most significantly enriched biological process ontologies with the largest aging-associated phase shifts are shown for male (blue) and female (orange) fibroblasts.

Bars represent the absolute change in phase (hours), with the corresponding FDR values displayed to the right of each bar. Pathways significantly enriched in young samples are shown in light colors (light blue for young males, light yellow for young females), those enriched in old samples appear in dark colors (dark blue for old males, dark orange for old females), and pathways enriched in both young and old are shown in grey.

**Supporting Tables**

**Table S1: Cosinor results of circadian clock genes from whole kidney RNA-Seq.** Cosinor results from all cohorts of circadian clock genes from whole kidney bulk RNA-Seq with both the raw and transformed data reported for each gene. With the raw model, the intercept, main_rrr1, and main_sss1 are reporting if the sine or cosine wave of expression is significant. For the transformed model, the intercept, amp1, and acr1 are reporting the amplitude and phase of the computed wave form.

**Table S2: DHARMa residual check for circadian clock genes in whole kidneys.** DHARMa residual results for all cohorts of circadian clock genes from whole kidney bulk RNA-Seq. The columns report all DHARMa computed measures with the model outcome stated in the DHARMa_any_fail column.

**Table S3: Cosinor results for senescence-associated genes from whole kidney RNA-Seq.** Cosinor results from all cohorts of senescence-associated genes from whole kidney bulk RNA-Seq with both the raw and transformed data reported for each gene. With the raw model, the intercept, main_rrr1, and main_sss1 are reporting if the sine or cosine wave of expression is significant. For the transformed model, the intercept, amp1, and acr1 are reporting the amplitude and phase of the computed wave form.

**Table S4: DHARMa residual check for senescence-associated genes in whole kidneys.** DHARMa residual results for all cohorts of senescence-associated genes from whole kidney bulk RNA-Seq. The columns report all DHARMa computed measures with the model outcome stated in the DHARMa_any_fail column.

**Table S5: Circadian genes in young male tail-tip fibroblasts.** ECHO results of circadian genes in young male tail tip fibroblasts.

**Table S6: Circadian genes in old male tail-tip fibroblasts.** ECHO results of circadian genes in old male tail tip fibroblasts.

**Table S7: Circadian genes in young female tail tip fibroblasts.** ECHO results of circadian genes in young female tail tip fibroblasts.

**Table S8: Circadian genes in old female tail tip fibroblasts.** ECHO results of circadian genes in old female tail tip fibroblasts.
